# MYC regulates ribosome biogenesis and mitochondrial gene expression programs through interaction with Host Cell Factor-1

**DOI:** 10.1101/2020.06.22.164764

**Authors:** Tessa M. Popay, Jing Wang, Clare M. Adams, Simona G. Codreanu, Stacy Sherrod, John A. McClean, Lance R. Thomas, Shelly L. Lorey, Yuichi J. Machida, April M. Weissmiller, Christine M. Eischen, Qi Liu, William. P. Tansey

**Author notes:** Corresponding author and contact: Department of Cell and Developmental Biology, Vanderbilt University School of Medicine, 465 21^st^ Avenue South, Nashville, TN 37232.

## Abstract

The oncoprotein transcription factor MYC is a major driver of malignancy and a highly-validated but challenging target for development of anti-cancer therapies. Novel strategies to inhibit MYC may come from understanding the co-factors it uses to drive pro-tumorigenic gene expression programs, providing their role in MYC activity is understood. Here, we interrogate how one MYC co-factor, Host Cell Factor (HCF)-1, contributes to MYC activity in a Burkitt lymphoma setting. We identify genes connected to mitochondrial function and ribosome biogenesis as direct MYC/HCF-1 targets, and demonstrate how modulation of the MYC–HCF-1 interaction influences cell growth, metabolite profiles, global gene expression patterns, and tumor growth *in vivo*. This work defines HCF-1 as a critical MYC co-factor, places the MYC–HCF-1 interaction in biological context, and highlights HCF-1 as a focal point for development of novel anti-MYC therapies.

## INTRODUCTION

MYC oncogenes (c-, L- and N-) encode a family of related transcription factors that are overexpressed in a majority of cancers and responsible for ~100,000 cancer-related deaths in the United States each year (Tansey, 2014). Capable of acting as both transcriptional activators and repressors, MYC proteins (hereafter ‘MYC’) dimerize with their obligate partner MAX (Blackwood and Eisenman, 1991) to bind to and regulate the expression of thousands of genes connected to the cell cycle, protein synthesis, metabolism, genome stability, apoptosis, and angiogenesis (Tansey, 2014). Fueled by reports that experimental inactivation of MYC promotes tumor regression in mice (Alimova et al., 2019; Beaulieu et al., 2019; Giuriato et al., 2006; Jain et al., 2002; Soucek et al., 2013), there is considerable interest in the idea that MYC inhibitors could form the basis of broadly-effective anti-cancer therapies. MYC itself, however, is widely viewed as undruggable (Dang et al., 2017), meaning that effective strategies to pharmacologically inhibit MYC will most likely come from targeting the co-factors it uses to drive and sustain the malignant state (Brockmann et al., 2013; Bryan et al., 2020).

Hundreds of proteins have been found to interact, directly or indirectly, with MYC (Baluapuri et al., 2020; Dingar et al., 2014; Kalkat et al., 2018; Kalkat et al., 2011). One effective strategy for prioritizing which of these potential co-factors to study has been to focus on those that interact with conserved segments of the MYC protein, which are referred to as “MYC boxes” (Mb) (Meyer and Penn, 2008). In addition to the highly-conserved basic helix-loop-helix domain that interacts with MAX, six evolutionarily conserved MYC boxes have been described (Baluapuri et al., 2020). On average, MYC boxes are around 15 amino acid residues in length, and although it is clear that they each mediate multiple protein-protein interactions (Kalkat et al., 2018), a number of predominant interactors have been described for most of these segments: Mb0, for example, interacts with the general transcription factor TFIIF to stimulate transcriptional elongation (Kalkat et al., 2018), MbI interacts with the ubiquitin ligase SCF^FBW7^ to control MYC protein stability (Welcker et al., 2004), MbII binds the STAGA component TRRAP (McMahon et al., 1998) to regulate histone acetylation (Kalkat et al., 2018), and MbIIIb interacts the chromatin-associated protein WDR5 (Thomas et al., 2015) to facilitate its recruitment to ribosomal protein genes (Thomas et al., 2019). The two remaining MYC boxes are less well understood, but MbIIIa is important for tumorigenesis (Herbst et al., 2005) and recruitment of HDAC3 to chromatin (Kurland and Tansey, 2008), and MbIV binds the ubiquitous chromatin-associated protein host cell factor (HCF)-1 (Thomas et al., 2016).

HCF-1 is an essential nuclear protein (Goto et al., 1997) that is synthesized as a 2035 amino acid precursor and proteolytically cleaved by O-GlcNAc transferase (OGT) (Capotosti et al., 2011) into amino-(HCF-1_N_) and carboxy-(HCF-1_C_) terminal segments that remain associated. HCF-1 was first identified through its ability to assemble into a multiprotein-DNA complex with the herpes simplex virus transactivator VP16, but was later shown to function in uninfected cells as a co-factor for cellular transcription factors, and as part of the Sin3 and MLL/SET histone modifying complexes (Wysocka and Herr, 2003). The interaction of HCF-1 with VP16 is direct and is mediated by a tetrapeptide “EHAY” motif in VP16 that binds to a region within the HCF-1_N_ fragment known as the VP16-induced complex (VIC) domain (Freiman and Herr, 1997). This four residue HCF-1-binding motif (HBM)—consensus (D/E)-H-x-Y—is present in other viral and cellular transcription factors that interact directly with the HCF-1 VIC domain, including key cell cycle regulators such as the E2F family of proteins (Tyagi et al., 2007).

We identified HCF-1 as a MYC-associated protein through proteomic approaches, and demonstrated that the interaction occurs through the VIC domain of HCF-1 and an atypical HBM within MbIV that carries the sequence ‘QHNY’ (Thomas et al., 2016). Mutation of these four HBM residues to alanine disrupts the interaction of MYC with HCF-1 *in vitro* and reduces the ability of MYC to promote murine fibroblast tumor growth in nude mice (Thomas et al., 2016).

The small and well-defined interaction point between MYC and HCF-1, and the importance of this interaction to tumorigenesis, raise the possibility that the MYC–HCF-1 nexus could be a viable venue for discovery of novel anti-MYC therapies. If that venue is to be pursued, however, we need to place this interaction in biological context, identify the gene networks that are under its control, and determine whether the MYC–HCF-1 interaction is required for tumor initiation, maintenance, or both. Here, we use a combination of loss- and gain-of function approaches to interrogate the role of the MYC–HCF-1 interaction in the context of a canonically MYC-driven cancer—Burkitt lymphoma. We demonstrate that the interaction between MYC and HCF-1 is directly involved in controlling the expression of genes linked to ribosome biogenesis, translation, and mitochondrial function. We define the impact of modulation of this interaction on cell growth, metabolism, and global gene expression patterns. And we show that disrupting the MYC–HCF-1 interaction promotes rapid and persistent tumor regression *in vivo*. This work reveals how MYC executes a core arm of its pro-tumorigenic gene expression changes, defines HCF-1 as a tumor-critical MYC co-factor, and provides proof-of-concept for a new way to inhibit MYC in the clinic.

## RESULTS

### Bidirectional modulation of the MYC−HCF-1 interaction

To understand the role of HCF-1 in MYC function, we sought to use separation-of-function mutations in MYC that modulate interaction with HCF-1 in a predictable way. We therefore introduced a number of mutations in the atypical HBM of MYC (QHNY) that we expected to decrease—or increase—interaction with HCF-1, based on properties of prototypical HBM sequences [(Freiman and Herr, 1997); Figure 1A]. Specifically, we substituted the MYC HBM for the canonical HBM from VP16 (VP16 HBM); we also mutated the invariant histidine of the HBM to glycine in the MYC (H307G) and VP16 (VP16 HBM:H307G) contexts, or we changed all four HBM residues to alanine (4A). A mutation in the separate WDR5-binding motif [WBM; (Thomas et al., 2015)] was our specificity control. We transiently-expressed these full-length FLAG-tagged MYC proteins in 293T cells and measured their ability to interact with endogenous HCF-1 in a co-immunoprecipitation (co-IP) assay (Figure 1B). As expected, the 4A mutation disrupts the MYC– HCF-1 interaction, as do both histidine to glycine substitutions—confirming the essentiality of this core HBM residue to the MYC–HCF-1 association. In contrast, replacing the MYC HBM with the canonical VP16 sequence increases the amount of HCF-1 recovered in the co-IP. This enhanced binding is likely a result of increasing the affinity of the direct MYC–HCF-1 interaction, as it is also observed with recombinant MYC and the HCF-1_VIC_ domain in *in vitro* binding assays (Figure 1 - figure supplement 1A). Based on these data, we conclude that the MYC HBM is an authentic and direct HCF-1 binding motif, and that its variation from the canonical HBM sequence leads to a tempered interaction with HCF-1. We also conclude that we can use the 4A and VP16 HBM mutations to probe the significance of the MYC–HCF-1 interaction through both loss- and gain-of-function approaches.

**Figure 1:**
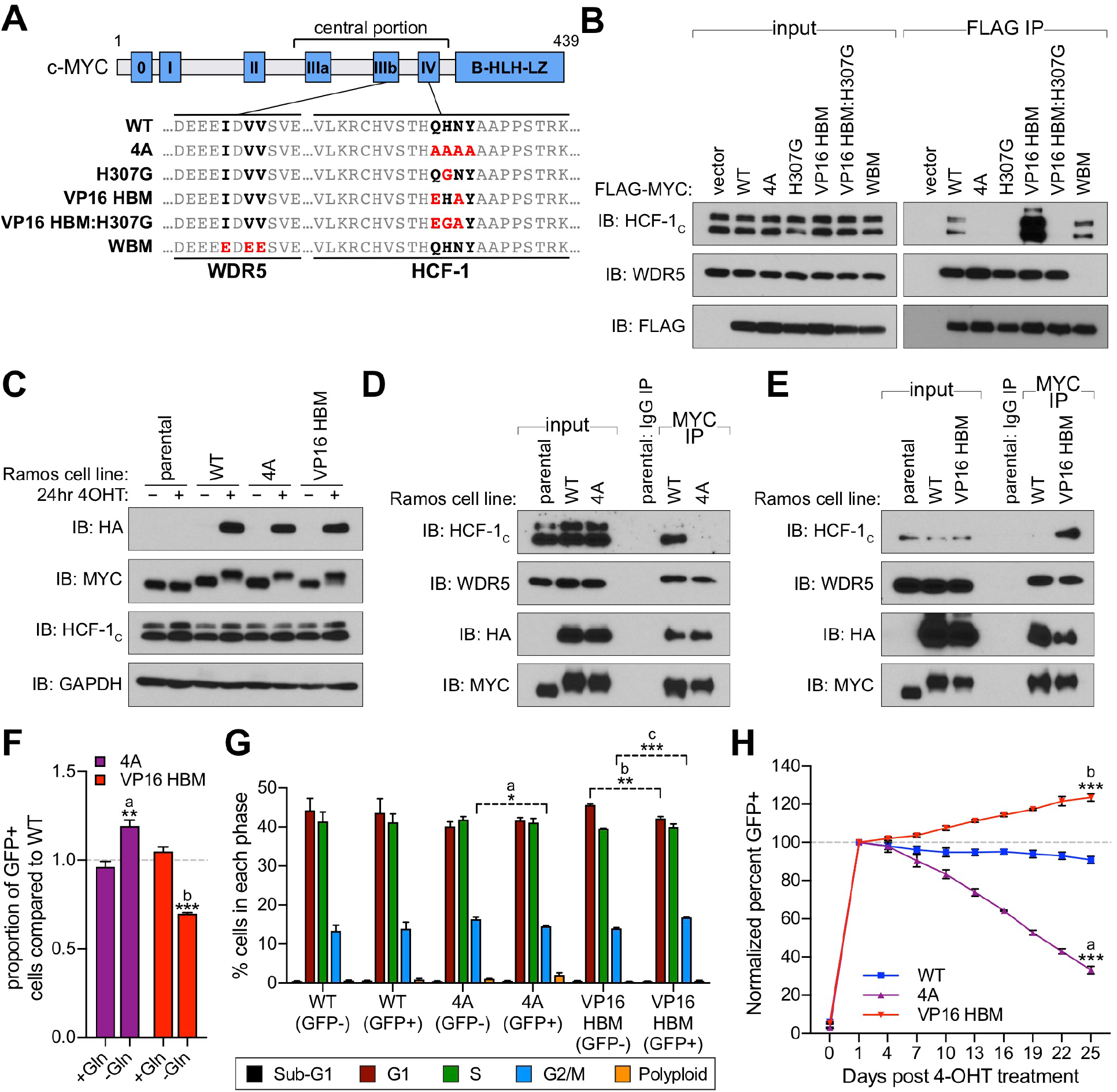
A gain- and loss-of-function system to study the MYC–HCF-1 interaction. (**A**) Schematic of MYC, depicting the location of the six MYC boxes (Mb0–MbIV). MbIIIb carries a WDR5-binding motif (WBM). MbIV contains an HCF-1 binding motif (HBM). Residues relevant to the WBM or HBM are in bold, and residues mutated in this study are in red. (**B**) FLAG-tagged full-length MYC proteins carrying the mutations described in (A) were transiently expressed in 293T cells, recovered by anti-FLAG immunoprecipitation (IP), and the input, or IP eluates, probed for the presence of HCF-1_C_, WDR5, and FLAG-tagged proteins by western blotting. (**C**) Western blot of lysates from parental (CRE-ER^T2^) or switchable Ramos cells (WT, 4A, or VP16 HBM) ± 20 nM 4-OHT for 24 hours. Blots were probed with antibodies against the HA tag, c-MYC, HCF-1_C_, and GAPDH. (**D**) and (**E**) Parental (CRE-ER^T2^) or switchable Ramos cells (WT, 4A, or VP16 HBM) were treated with 20 nM 4-OHT for 24 hours, lysates prepared, and IP performed using anti-IgG or anti-MYC antibodies. Input lysates and IP eluates were probed using antibodies against HCF-1_C_, WDR5, HA tag, and MYC by western blotting. (**F**) Switchable Ramos cell lines were pulsed with 20 nM 4-OHT for two hours to switch approximately 50% of cells, propagated for three days, and grown for 16 hours in media with or without glutamine. The impact of glutamine deprivation was measured by flow cytometry to determine the proportion of GFP-positive (switched) cells. For each of the mutants, the proportion of GFP-positive cells was normalized to that for WT cells. Shown are the mean and standard error for three biological replicates. Student’s t-test between +Gln and -Gln was used to calculate P-values; a = 0.0066, b = 0.0002. (**G**) Switchable Ramos cells were pulsed with 4-OHT as in (F), grown for seven days, and cell cycle distribution determined by propidium iodide (PI) staining and flow cytometry, binning cells according to whether they expressed GFP (GFP+, switched) or not (GFP−, unswitched). Shown are the mean and standard error for three biological replicates. Student’s t-test between GFP− and GFP+ cells was used to calculate P-values; a = 0.033, b = 0.0041, c = 0.0006. (**H**) Switchable Ramos cells were pulsed with 4-OHT as in (F), and the proportion of GFP-positive cells measured by flow cytometry 24 hours after treatment and every three days following. For each of the replicates, the proportion of GFP-positive cells is normalized to that on day one. Shown are the mean and standard error for three biological replicates. Sudent’s t-test between WT and each of the mutants at day 25 was used to calculate P-values; a=0.000028, b=0.00026. The following figure supplement is available for figure 1: **Figure 1 - figure supplement 1**: Validation of switchable Ramos cell lines to study the MYC–HCF-1 interaction.

### The MYC–HCF-1 interaction stimulates proliferation of Burkitt lymphoma cells

To understand the cellular consequences of modulating the MYC–HCF-1 interaction, we engineered a system that allows us to express the 4A or VP16 HBM mutant MYC proteins as the sole form of MYC in a cell. We chose Ramos cells, a Burkitt lymphoma (BL)-derived line in which a *t(8;14)* translocation places one *c-MYC* allele under regulatory control of the immunoglobulin heavy chain enhancer (Figure 1 - figure supplement 1B) (Wiman et al., 1984). The untranslocated *c-MYC* allele is not expressed in these cells (Bemark and Neuberger, 2000). Because sequences encoding the MYC HBM are contained within exon 3, we used CRISPR/Cas9-triggered homologous recombination of the translocated *MYC* allele to integrate an exon 3 switchable cassette for wild-type (WT) MYC, 4A, or VP16 HBM mutants, and confirmed appropriate integration by Southern blotting (Figure 1 - figure supplement 1C and 1D). In cells expressing an inducible CRE-ER^T2^ recombinase, treatment with 4-hydroxytamoxifen (4-OHT) results in the excision of exon 3 of *MYC*, bringing in place a modified exon 3 that carries an HA-epitope tag and drives expression of P2A-linked green fluorescent protein (GFP). Twenty four hours after 4-OHT treatment, at least 85% of cells in each population are switched—as monitored by GFP expression (Figure 1 - figure supplement 1E), and we observe the expected appearance of HA-tagged MYC proteins, which migrate more slowly due to the presence of the epitope tag (Figure 1C). Importantly, the exchanged MYC proteins are expressed at levels comparable to endogenous MYC (Figure 1C), and behave as expected, with the 4A mutant showing reduced (Figure 1D), and the VP16 HBM mutant enhanced (Figure 1E), interaction with endogenous HCF-1. Also as expected, these mutations have minimal impact on the interaction of MYC with WDR5. Thus, we successfully generated a system for inducible, selective, and bidirectional modulation of the MYC−HCF-1 interaction in the context of an archetypal MYC-driven cancer cell line.

To monitor the contribution of the MYC–HCF-1 interaction to cell proliferation, we pulsed each of our engineered Ramos lines with 4-OHT for two hours to generate approximately equally mixed populations of switched and unswitched cells. We then monitored how the GFP-positive switched cells in the population compared to their unswitched counterparts in terms of glutamine-dependency (Figure 1F), cell cycle profiles (Figure 1G), and long-term growth (Figure 1H). We see that 4A switched cells have a selective advantage over the WT switch in their ability to grow without exogenous glutamine (Figure 1F). This advantage is likely due to loss of the MYC–HCF-1 interaction, as the VP16 HBM mutant cells have a corresponding deficit in growth under glutamine-starvation conditions (Figure 1F). Compared to their wild-type counterparts, cell cycle profiles for the two mutants are not dramatically altered, but we did observe small but statistically significant changes in the proportion of cells in G_2_/M (Figure 1G), which again trend in opposite directions for the two MYC mutants—decreasing for the 4A-expressing cells and increasing for those that express the VP16 HBM mutant (Figure 1G). And finally, by monitoring long-term growth, we observe that 4A mutant cells are gradually lost from the culture over time, whereas there is a significant enrichment of VP16 HBM cells, compared to the WT control switch (Figure 1H). The altered and opposing impact of the 4A and VP16 HBM mutations in these assays leads us to conclude that the MYC–HCF-1 interaction promotes the glutamine-dependency—and rapid proliferative status—of these BL cells in culture.

### The MYC–HCF-1 interaction influences intracellular amino acid levels

As part of our survey of the impact of the MYC–HCF-1 interaction on Ramos cell processes, we determined whether metabolite levels are altered in response to expression of the 4A or VP16 HBM MYC mutants. We performed global, untargeted, mass spectrometry-based metabolomics on switched cells using reverse-phase liquid chromatography (RPLC) and hydrophilic interaction liquid chromatography (HILIC) separation methods, detecting ~2,000 metabolites with each approach (Figure 2A–F). In general, more metabolites are significantly changed for the 4A than the VP16 HBM MYC mutant (Figure 2 - figure supplements 1A and 1B, Figure 2 - Source Data 1 and 2), and the magnitude of changes are greater for the 4A protein. Significantly-changed metabolites group into a variety of categories, with the majority being amino acid-related or lipid-based (Figure 2C and 2F). For the 4A mutant, we observe significant enrichment in pathways connected to nitrogen and glycerophospholipid metabolism, as well as those associated with amino acid metabolism and utilization, including alanine, aspartate, and glutamate metabolism (Figure 2G). Similar trends are apparent for the VP16 HBM mutant, but pathway analysis failed to reach statistical significance, likely due to the smaller number of altered metabolites in these cells.

**Figure 2:**
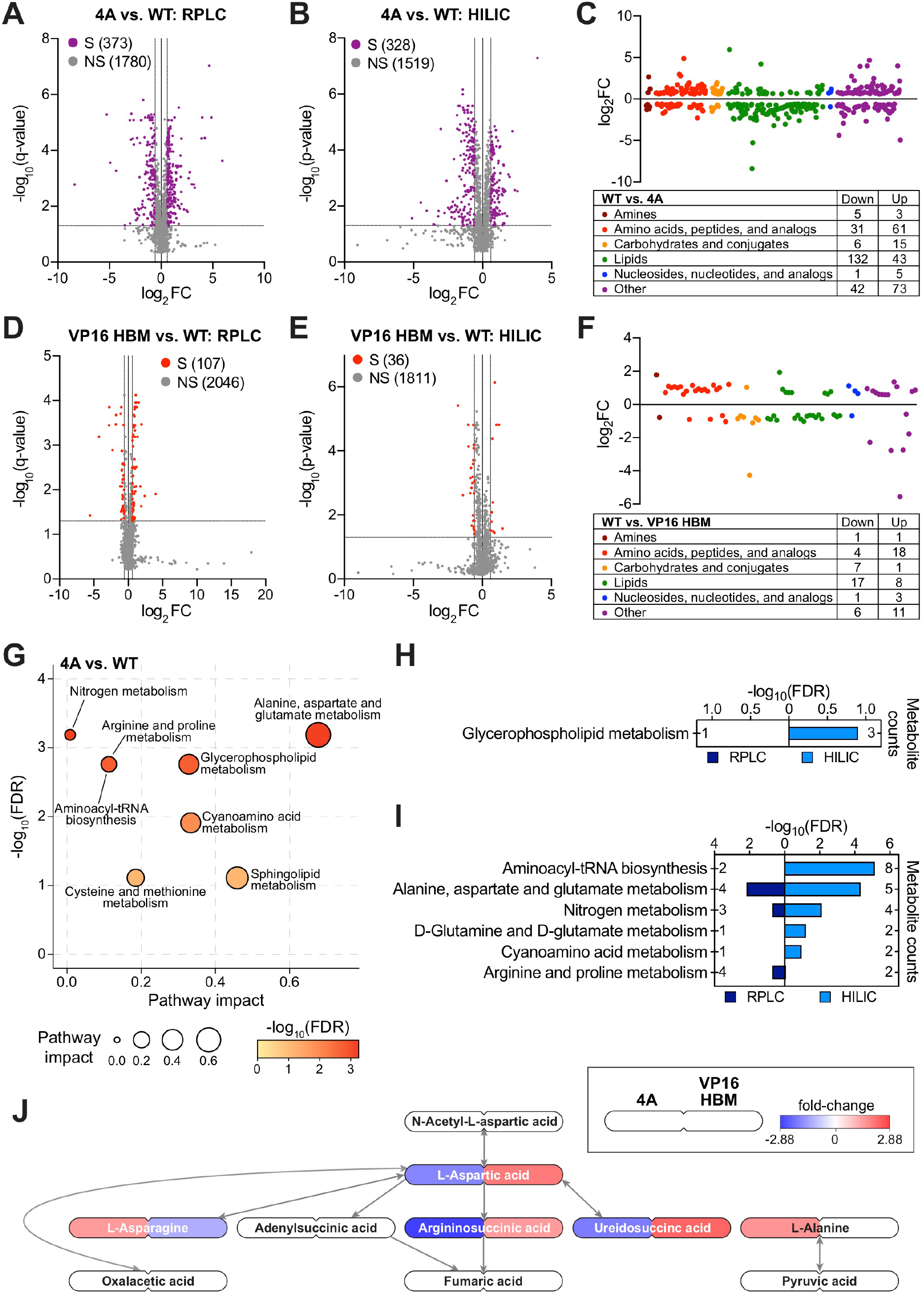
The MYC–HCF-1 interaction influences intracellular amino acid levels. (**A**) and (**B**) Volcano plots of metabolites detected by RPLC (A) or HILIC (B) in WT and 4A switchable Ramos cells treated for 24 hours with 20 nM 4-OHT. Metabolites that were significantly (S) changed (false discovery rate, FDR < 0.05 & |FC| > 1.5) with the 4A MYC mutant compared to WT are colored. Non-significant (NS) changes are in grey. Five biological replicates for WT and four biological replicates for 4A were used to calculate FDR and fold-changes (FC). (**C**) Classification of metabolites that were significantly changed (FDR < 0.05 & |FC| > 1.5) with the 4A mutant compared to WT cells. (**D**) and (**E**) Volcano plots of metabolites detected by RPLC (D) or HILIC (E) in WT and VP16 HBM switchable Ramos cells treated for 24 hours with 20 nM 4-OHT. Metabolites that were significantly (S) changed (FDR < 0.05 & |FC| > 1.5) with the VP16 HBM MYC mutant compared to WT are colored. Non-significant (NS) changes are in grey. Five biological replicates for WT and VP16 HBM were used to calculate FDR and fold-changes. (**F**) Classification of metabolites that were significantly changed (FDR < 0.05 & |FC| > 1.5) with the VP16 HBM mutant compared to WT cells. (**G**) Enrichment analysis of KEGG pathways using a combined list of significantly changed metabolites (FDR < 0.05 & |FC| > 1.5) with the 4A mutant from HILIC and RPLC. Pathway impact reflects centrality and the number of matched metabolites. Pathways with FDR < 0.2 are shown. (**H**) Clusters of annotated metabolites from Figure 2 - figure supplement 1C that were changed in the same direction for the 4A and VP16 HBM mutants were independently subjected to pathway enrichment analysis. Pathways with FDR < 0.2 for either RPLC and HILIC are shown. (**I**) As in (H), except for of annotated metabolites from Figure 2 - figure supplement 1C that were changed in opposite directions in the 4A and VP16 HBM mutants. (**J**) Metabolites (FDR < 0.05) in the “alanine, separtate, and glutamate metabolism” pathway that were impacted by the 4A (left) and VP16 HBM (right) MYC mutants. Node color represents the fold-change over WT. The remainder of pathway is shown in **Figure 2 - figure supplement 1F and 1G**. The following source data and figure supplement(s) are available for figure 2: **Figure 2 - Source Data 1:** Ramos 4A and VP16 HBM RPLC significant changes **Figure 2 - Source Data 2:** Ramos 4A and VP16 HBM HILIC significant changes **Figure 2 - figure supplement 1:** Impact of the 4A and VP16 HBM mutants on Ramos cell metabolism.

Comparing the direction of individual metabolite changes for the 4A and VP16 HBM mutants (Figure 2 - figure supplement 1C), we note that a significant portion of the metabolite changes detected by both the RPLC and HILIC methods are in the same direction for the two MYC mutants. In general, these shared metabolite changes fail to cluster strongly into biological pathways; the only significantly enrichment being glycerophospholipid metabolism (Figure 2H). Focusing on metabolite changes that occur in opposite directions for the 4A and VP16 HBM mutants, however, we observe significant enrichment in pathways linked to nitrogen and amino acid metabolism (Figure 2I). There is a clear anti-correlation between the impact of the 4A and VP16 HBM mutations on metabolites connected to aspartic acid (Figure 2J), and we observe that intracellular levels of glutamine (and associated metabolites) are increased in the 4A, and decreased in the VP16 HBM mutant cells (Figure 2 - figure supplement 1D and 1E). Notably, these changes in intracellular amino acid levels are not confined to aspartic acid and glutamine, but rather there is a general tendency for amino acid levels to be increased in 4A and decreased in VP16 HBM mutant cells, compared to the WT switch (Table 1). Based on these data, we conclude that the MYC–HCF-1 interaction, directly or indirectly, plays a global role in influencing intracellular amino acid levels in this setting.

**Table 1:** Modulating the MYC–HCF-1 interaction modulates intracellular amino acid levels. All data are derived from itchable Ramos cells treated with 20 nM 4-OHT for 24 hours. Amino acid levels were measured following separations HILIC. FDR and fold-changes (FC) were calculated between the mutants and WT. Five biological replicates for WT and P16 HBM and four biological replicates for 4A were analyzed. FDR < 0.05 are highlighted in green, FC > 0 in red, and FC < 0 in blue. Confidence levels reflect the confidence in metabolite identification; L1 is validated, L2 is putative, and L3 tentative. ND=not detected; NS=not significant.

|  | 4A vs. WT |  | VP16 HBM vs. WT |  | Confidence level |
| --- | --- | --- | --- | --- | --- |
|  | FDR | FC | FDR | FC |  |
| **Glycine** | 2.52E-04 | 1.31 | 3.11E-05 | -1.48 | L3 |
| **L-Alanine** | 9.97E-05 | 1.62 | NS |  | L2 |
| **L-Arginine** | 2.31E-02 | 1.18 | NS |  | L1 |
| **L-Asparagine** | 6.67E-05 | 1.50 | 2.02E-03 | -1.23 | L2 |
| **L-Aspartic acid** | 1.24E-03 | -1.37 | 1.54E-05 | 2.01 | L2 |
| **L-Cysteine** | ND |  | ND |  |  |
| **L-Glutamic acid** | 2.53E-03 | 1.46 | NS |  | L2 |
| **L-Glutamine** | 2.33E-05 | 1.46 | 9.38E-05 | -1.41 | L2 |
| **L-Histidine** | 8.70E-05 | 1.64 | NS |  | L2 |
| **L-Isoleucine** | 1.10E-03 | 1.39 | 2.36E-02 | -1.21 | L2 |
| **L-Leucine** | 8.63E-04 | 1.33 | 2.55E-02 | -1.18 | L2 |
| **L-Lysine** | 1.66E-02 | 1.19 | NS |  | L2 |
| **L-Methionine** | 6.51E-05 | 1.61 | 4.39E-03 | -1.27 | L2 |
| **L-Phenylalanine** | 4.71E-04 | 1.48 | 3.46E-02 | -1.22 | L2 |
| **L-Proline** | 1.01E-02 | 1.17 | NS |  | L2 |
| **L-Serine** | 7.48E-05 | 1.59 | NS |  | L1 |
| **L-Threonine** | 9.63E-05 | 1.54 | 1.25E-02 | -1.17 | L2 |
| **L-Tryptophan** | 1.56E-04 | 1.64 | NS |  | L1 |
| **L-Tyrosine** | 1.22E-04 | 1.59 | 1.9E-02 | -1.18 | L2 |
| **L-Valine** | 4.41E-02 | 1.14 | NS |  | L3 |

### The MYC–HCF-1 interaction influences expression of genes connected to ribosome biogenesis and the mitochondrial matrix

Next, we used RNA-sequencing (RNA-Seq) to monitor transcriptomic changes associated with modulating the MYC–HCF-1 interaction. Twenty four hours after switching, we observed changes in the levels of ~4,000 transcripts in the 4A, and ~3,600 transcripts in the VP16 HBM, cells compared to the WT switch (Figure 3 - figure supplement 1A and 1B, Figure 3 - Source Data 1 and 2). For both mutants, these changes are modest in magnitude (Figure 3A), consistent with what is typically reported for MYC (Levens, 2002; Nie et al., 2012). Gene ontology (GO) enrichment analysis revealed that transcripts decreased by introduction of the 4A mutant are strongly linked to ribosome biogenesis, tRNA metabolism, and the mitochondrial matrix (Figure 3B), while those that are induced have links to transcription, cholesterol biosynthesis, and chromatin. For the VP16 HBM MYC mutant, decreased transcripts cluster in categories related to the centrosome and the cell cycle (Figure 3C). What is particularly striking, however, is that genes that are induced by the VP16 HBM protein have a pattern of clustering that is almost the exact opposite of those suppressed by the 4A mutant—including ribosome biogenesis, tRNA metabolism, and the mitochondrial matrix.

**Figure 3:**
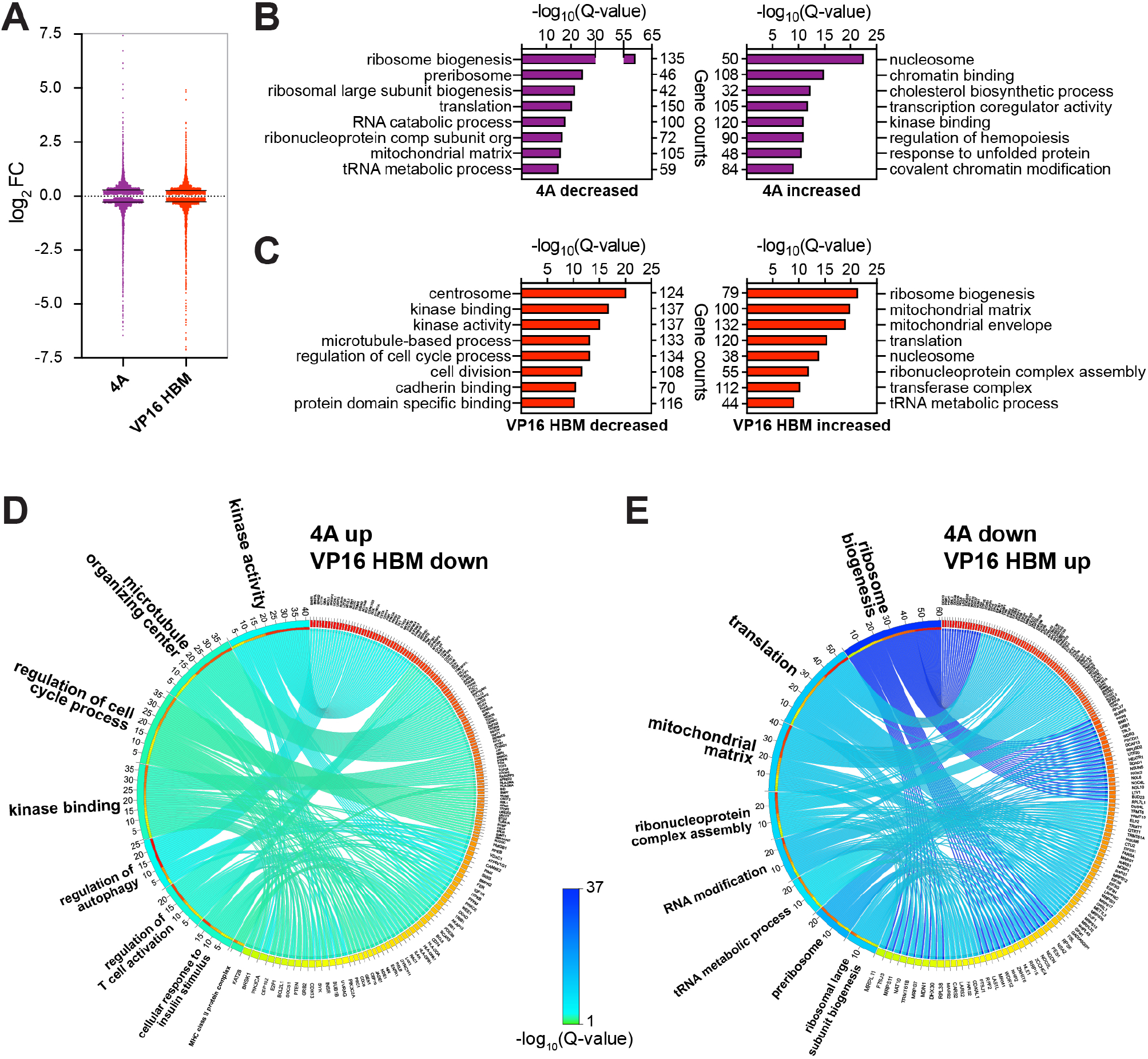
The MYC–HCF-1 interaction influences the expression of genes involved in ribosome biogenesis and mitochondrial pathways. Switchable Ramos cells were treated with 20 nM 4-OHT for 24 hours, RNA isolated, and RNA-Seq performed. (**A**) Scatter plot showing the distribution of log_2_FC of significant (FDR < 0.05) RNA-Seq changes with the 4A and VP16 HBM MYC mutants, compared to the WT switch. Solid lines represents the median log_2_FC for decreased (4A: −0.2858; VP: −0.2747) and increased (4A: 0.281; VP: 0.2558) genes compared to WT. For clarity, some data points were excluded; these data points are highlighted in Figure 3 - Source Data 1 and 2. (**B**) and (**C**) Categories from the top eight families in GO enrichment analysis of significant (FDR < 0.05) gene expression changes under each condition (B: 4A; C:VP16 HBM). (**D**) and (**E**) Categories from the top eight families in GO enrichment analysis of the anti-correlative gene clusters shown in Figure 3 - figure supplement 1C, for genes that were decreased (D) or increased (E) with the 4A mutant. The Q-value of categories is represented by the ribbon color, which is scaled across these figures and **Figure 3 - figure supplement 1E and 1F**. Categories are ranked by the number of matched genes, and genes are ranked by the number of categories in which they fall. The following source data and figure supplement(s) are available for figure 3: **Figure 3 - Source Data 1:** Ramos 4A 24 hour RNA-Seq significant changes **Figure 3 - Source Data 2:** Ramos VP16 HBM 24 hour RNA-Seq significant changes **Figure 3 - Source Data 3:** Ramos 4A and VP16 HBM 24 hour RNA-Seq shared significant changes **Figure 3 - figure supplement 1:** Gene expression changes induced by the 4A and VP16 HBM mutants. **Figure 3 - figure supplement 2:** Amino acids and their cognate tRNA-ligases are influenced by the MYC–HCF-1 interaction.

If anti-correlations between these gain- and loss-of-function mutants can be used to reveal MYC–HCF-1 co-regulated processes, the above data highlight protein synthesis and mitochondrial function as key points of convergence for the interaction of MYC with HCF-1. To explore this on a gene-by-gene basis, we compared individual gene expression changes that were either the same, or opposite, in direction for the 4A and VP16 HBM mutants (Figure 3 - figure supplement 1C and 1D, Figure 3 - Source Data 3). Transcripts decreased by both mutations show modest enrichment in categories connected to immune signaling and cell adhesion (Figure 3 - figure supplement 1E), whereas increased transcripts are primarily enriched in those encoding histones (Figure 3 - figure supplement 1F). Turning to transcripts that change in opposite directions with each mutant, those that are induced by the 4A mutation are moderately enriched in categories relating to kinase function and the cell cycle (Figure 3D), while those that are reduced by the 4A mutant are strongly enriched in categories connected to ribosome biogenesis and the mitochondrial matrix (Figure 3E). This analysis confirms that reciprocal changes we observed for the GO categories in Figure 3B and 3C results from reciprocal changes in the expression of a common set of genes. From our data, we conclude that the MYC–HCF-1 interaction plays an important role in influencing the expression of genes that promote ribosome biogenesis and maintain mitochondrial function.

Finally, we interrogated our RNA-Seq dataset for transcript changes that would correlate with the widespread changes in amino acid levels that occur upon modulation of the MYC–HCF-1 interaction. Here, we discovered that the accumulation of amino acids we observe with the 4A mutant is generally matched with a decrease in transcription of cognate aminoacyl-tRNA synthetases (Figure 3 - figure supplement 2A)—and vice-versa for the decreased amino acid levels in the gain-of-function VP16 HBM mutant (Figure 3 - figure supplement 2B). The reciprocal way in which amino acid levels and tRNA ligase expression changes in response to the 4A and VP16 HBM mutants is consistent with the idea that defects in tRNA charging lead to compensatory changes in amino acid uptake (Guan et al., 2014; Harding et al., 2000), and further reinforces the concept that a key biological context in which MYC and HCF-1 function together is protein synthesis.

### Ribosome biogenesis and mitochondrial matrix genes respond rapidly to HCF-1 depletion

As a challenge to the concept that ribosome biogenesis and mitochondrial matrix genes are controlled via the MYC–HCF-1 interaction, we asked whether expression of these genes is impacted by acute depletion of HCF-1, mediated via the dTAG method (Nabet et al., 2018). We used CRISPR/Cas9-triggered homologous recombination to integrate an mCherry-P2A-FLAG-FKBP12^F36V^ cassette into the *HCFC1* locus in Ramos cells; the effect of which is to amino-terminally tag HCF-1_N_ with the FLAG epitope and FKBP12^F36V^ tags, and to mark the population of modified cells by mCherry expression (Figure 4 - figure supplement 1A). Because the *HCFC1* locus resides on the X-chromosome, and because Ramos cells are derived from an XY patient, only a single integration event is needed. Tagged cells sorted by fluorescence-activated cell sorting display the expected shift in apparent molecular weight of HCF-1_N_ and the appearance of an appropriately-sized FLAG-tagged species (Figure 4A). Addition of the dTAG-47 degrader results in the rapid and selective disappearance of the HCF-1_N_ fragment; the HCF-1_C_ fragment is largely unaffected by up to 24 hours of dTAG-47 treatment (Figure 4B). Consistent with the known functions of HCF-1 (Julien and Herr, 2003), treated cells display altered cell cycle profiles (Figure 4 - figure supplement 1B), but appear to be able to complete at least one round of cell division, as notable deficits in proliferation are only evident 48 hours after dTAG-47 addition (Figure 4C). These data reveal that the HCF-1_N_ fragment is essential in Ramos cells, and that early analyses should be resistant to complicating effects of HCF-1_N_ degradation on cell proliferation.

**Figure 4:**
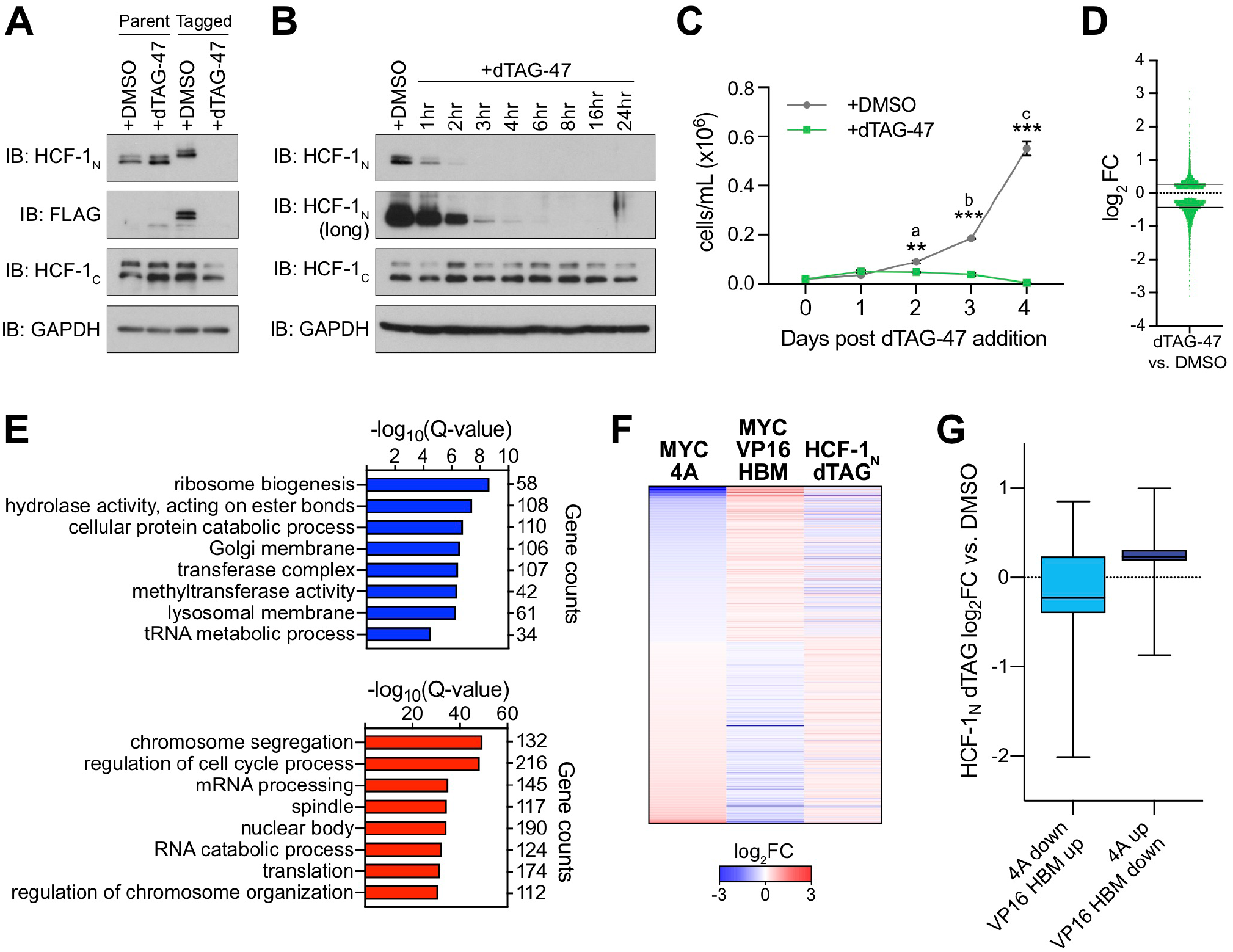
Genes regulated by the MYC–HCF-1 interaction are impacted by loss of HCF-1. (**A**) Western blot, comparing effect of treating untagged, parental cells or FKBP^FV^-HCF-1_N_ Ramos cells with DMSO or 500 nM dTAG-47 for 24 hours. Blots for HCF-1_N_, FLAG tag, HCF-1_C_, and GAPDH are shown. (**B**) Western blot of lysates from FKBP^FV^-HCF-1_N_ Ramos cells treated with 500 nM dTAG-47 for varying times, compared to cells treated with DMSO for 24 hours. Shown are short and long exposures of HCF-1_N_, and HCF-1_C_, with a GAPDH loading control. (**C**) Growth curve of FKBP^FV^-HCF-1_N_ Ramos cells treated with DMSO or 500 nM dTAG-47. Cells were counted every 24 hours for four days after plating. Shown are the mean and standard error for three biological replicates. Student’s t-test was used to calculate P-values; a = 0.0029, b = 0.000051, c = 0.000040. (**D**) Scatter plot showing the distribution of log_2_FC in RNA-Seq comparing DMSO to three hours of 500 nM dTAG-47 treatment (degradation of HCF-1_N_). Solid lines represent the median log_2_FC for decreased (−0.425655) and increased (0.270428) genes. For clarity, one data point was excluded; this data point is highlighted in Figure 4 - Source Data 1. (**E**) GO enrichment analysis of genes significantly (FDR < 0.05) decreased (top) and increased (bottom) in expression following treatment of FKBP^FV^-HCF-1_N_ Ramos cells with dTAG-47 for three hours. Excluded from this analysis are genes that were significantly changed when parental Ramos cells were treated with dTAG-47 for three hours. (**F**) Heatmap showing the log_2_FC of genes with significantly (FDR < 0.05) changed expression, as measured by RNA-Seq, under all conditions (switchable MYC alleles, 4A and VP16 HBM, and FKBP^FV^-HCF-1_N_ Ramos cells). Genes are clustered according to the relationship in expression changes between the 4A and VP16 HBM mutants, and ranked by the log_2_FC for the 4A mutant. Scale of heatmap is limited to [−3,3]. (**G**) Box-and-whisker plot showing the relationship between genes that are anti-correlated between the 4A and VP16 HBM MYC mutants, and significantly changed with the degradation of HCF-1_N_. Box denotes the 25th to 75th percentile, middle line marks the median, and whiskers extend from minimum to maximum point. The following source data and figure supplement(s) are available for figure 4: **Figure 4 - Source Data 1:** Ramos HCF-1_N_ degradation RNA-Seq significant changes **Figure 4 - Source Data 2:** Ramos untagged RNA-Seq significant changes **Figure 4 - figure supplement 1:** Inducible degradation of HCF-1_N_.

We performed RNA-Seq analysis three hours after addition of dTAG-47—a time point at which the majority of HFC-1_N_ is degraded (Figure 4B). Despite the early timepoint, we identified ~4,500 significant transcriptional changes associated with dTAG-47 treatment of sorted cells (Figure 4 - figure supplement 1C, Figure 4 - Source Data 1). These changes are equally divided between increased and decreased, although decreased transcripts are generally more impacted (larger median fold-change) than those that are induced (Figure 4D). Seventy-five of these differentially expressed genes are also altered in response to dTAG-47 treatment of untagged Ramos cells (Figure 4 - figure supplement 1D and 1E, Figure 4 - Source Data 2), and were excluded from further analyses. GO enrichment analysis showed that transcripts reduced by HCF-1_N_ degradation are similar in kind to those reduced by the 4A mutation in MYC—including ribosome biogenesis and tRNA metabolic processes (Figure 4E)—while those induced by HCF-1_N_ degradation tend to be cell cycle-connected (Figure 4E and Figure 4 - figure supplement 1F). Importantly, many of the genes that are differentially expressed upon HCF-1_N_ degradation are differentially expressed in the presence of either the 4A or VP16 HBM mutants (Figure 4 - figure supplement 1G), and we identified a union set of ~450 genes—oppositely regulated by the 4A and VP16 HBM mutants—the expression of which also changes when HCF-1_N_ is destroyed (Figure 4F). Within this set, loss of HCF-1_N_ tends to mimic the loss of function 4A mutant (Figure 4F), particularly for transcripts that are reduced when the MYC–HCF-1 interaction is disrupted (Figure 4G). Moreover, within the cohort of transcripts that are reduced by both HCF-1_N_ destruction and the 4A mutation, we see clear representation of genes connected to ribosome biogenesis and the mitochondrial matrix (Figure 4 - figure supplement 1H and 1I). Although performing RNA-Seq at this early time underestimates the impact of loss of HCF-1_N_ on the transcriptome, the presence of these ribosome biogenesis and mitochondrial matrix genes at the point of coalescence of all our RNA-Seq experiments strongly suggests that they are directly controlled the MYC–HCF-1 interaction.

### Most HCF-1 binding sites on chromatin are bound by MYC

To help identify direct transcriptional targets of the MYC–HCF-1 interaction, we next compared the genomic locations bound by MYC and HCF-1_N_ in Ramos cells. We performed ChIP-Seq using an antibody against the amino-terminus of HCF-1 (Machida et al., 2009), and identified ~1,900 peaks for HCF-1_N_ (Figure 5 - Source Data 1), the majority of which are promoter proximal (Figure 5A). These peaks occur at genes enriched in functions connected to HCF-1 (Minocha et al., 2019), including the mitochondrial envelope, the cell cycle, as well as ribonucleoprotein complex biogenesis (Figure 5B). Known (Figure 5C) and *de novo* (Figure 5 - figure supplement 1A) motif analysis revealed that HCF-1_N_ peaks are enriched in DNA sequences linked to nuclear respiratory factor (NRF)-1, as well as the Sp1/Sp2 family of transcription factors. Interestingly, although HCF-1 has not previously been linked directly to NRF-1 or this element, the motif is also a functional, non-canonical, E-box that MYC proteins are known to bind (Blackwell et al., 1993; Morrish et al., 2003). Consistent with enrichment of this variant E-box element in the HCF-1_N_ peaks, overlaying these data with our previous ChIP-Seq analysis of MYC in Ramos cells (Thomas et al., 2019) (Figure 5 - figure supplement 1B), we see that 85% of these HCF-1_N_ peaks are also bound by MYC (Figure 5D, Figure 5 - Source Data 2). The relationship between MYC and HCF-1_N_ at these sites is intimate (Figure 5E and Figure 5 - figure supplement 1C), and sites of co-binding tend to have higher signals for MYC (Figure 5F) and HCF-1 (Figure 5 - figure supplement 1D) than instances where each protein binds alone. As expected from the strong coalescence of MYC and HCF-1_N_ binding events, the properties of shared MYC–HCF-1_N_ peaks are very similar to those of HCF-1_N_ alone, in terms of location (Figure 5 - figure supplement 1E), GO enrichment categories (Figure 5 - figure supplement 1F), and motif representation (Figure 5 - figure supplement 1G). We conclude that most HCF-1_N_ binding sites on chromatin in Ramos cells occur at promoter proximal sites and that the majority of these sites are also bound by MYC.

**Figure 5:**
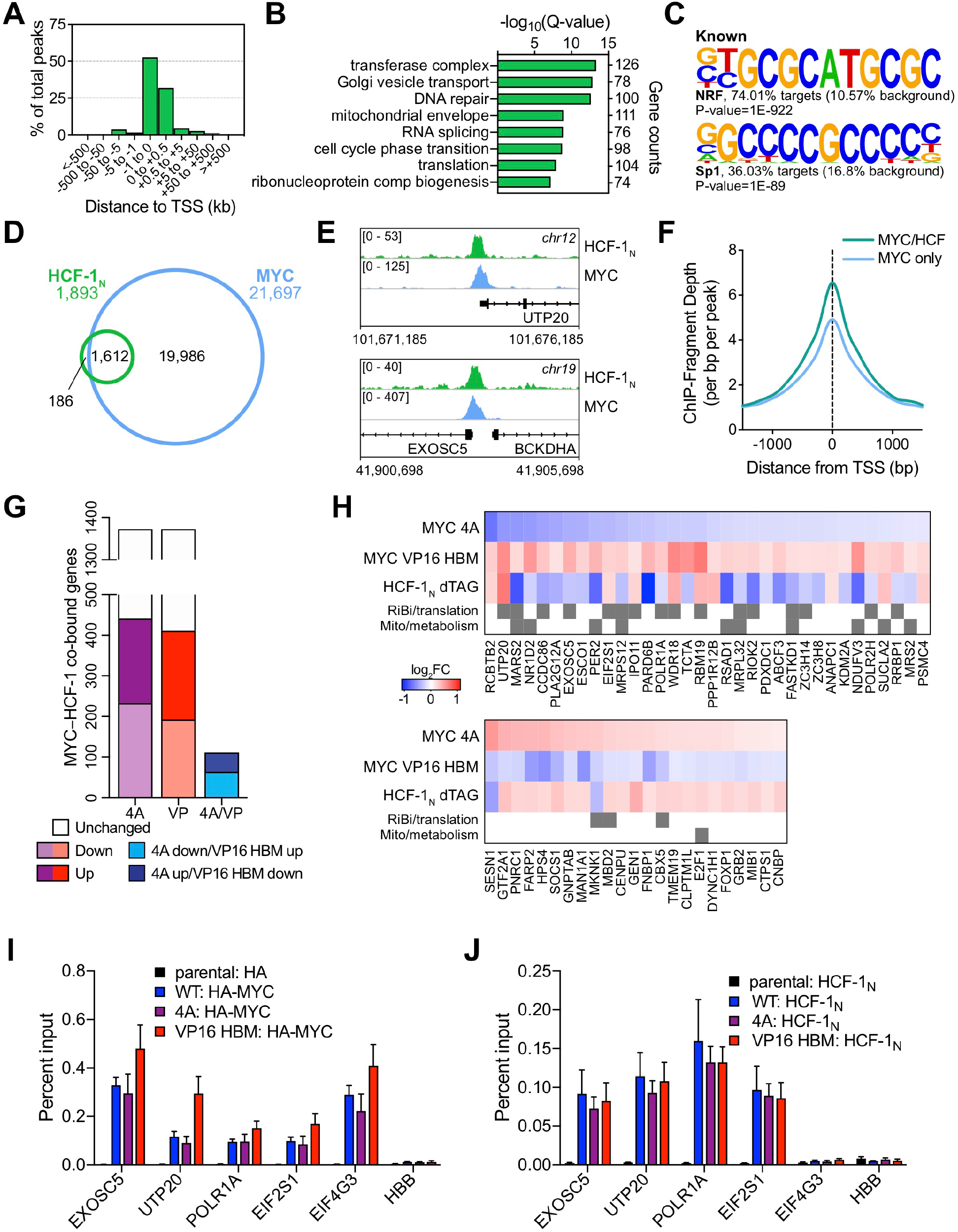
MYC is a widespread binding partner of HCF-1 on chromatin. (**A**) Distribution of HCF-1_N_ peaks in Ramos cells in relation to the nearest TSS, as determined by ChIP-Seq. (**B**) GO categories strongly represented in genes nearest to HCF-1_N_ peaks in Ramos cells. (**C**) Known motif analysis of HCF-1_N_ peaks in Ramos cells. Two of the most highly enriched motifs are shown, as well the percentage of target and background sequences with the motif, and the P-value. (**D**) Venn diagram showing HCF-1_N_ and MYC peaks in Ramos cells, and the number of regions that overlap between the datasets. ChIP-Seq data for MYC are from GSE126207. (**E**) Example Integrative Genomics Viewer (IGV) screenshots of regions that have overlapping peaks for MYC and HCF-1_N_ in Ramos cells. (**F**) Normalized MYC ChIP-Seq fragment counts where peaks overlap with HCF-1_N_ (MYC/HCF), compared to where they do not overlap (MYC) in Ramos cells. Data are smoothed with a cubic spline transformation. (**G**) Relationship between protein-coding genes that are co-bound by promoter-proximal MYC and HCF-1_N_ by ChIP-Seq, and are significantly (FDR < 0.05) decreased or increased in response to the 4A or VP16 HBM mutations. Also shown are genes where the expression is anti-correlated between the 4A and VP16 HBM mutants. (**H**) Heatmap showing genes that are co-bound by promoter proximal MYC and HCF-1_N_ in Ramos cells, have anti-correlative gene expression changes between for the 4A and VP16 MYC mutants, and have significant gene expression changes with HCF-1_N_ degradation. Genes that fall into GO categories relating to ribosome biogenesis or translation (RiBi/translation), and mitochondrial function or metabolism (Mito/metabolism) are highlighted. (**I**) ChIP, using anti-HA antibody, was performed on parental or switchable Ramos cells treated for 24 hour with 20 nM 4-OHT. Enrichment of genomic DNA was monitored by qPCR using primers that amplify across peaks. *HBB* is a negative locus for HA-MYC. ChIP efficiency was measured based on the percent recovery from input DNA. Shown are the mean and standard error for three biological replicates. (**J**) ChIP, using anti-HCF-1_N_ antibody, was performed on parental or itchable Ramos cells treated for 24 hour with 20 nM 4-OHT. Enrichment of genomic DNA was monitored by qPCR ing primers that amplify across peaks. *EIF4G3* and *HBB* are negative loci for HCF-1_N_. ChIP efficiency was measured based on the percent recovery from input DNA. Shown are the mean and standard error for three biological replicates. The following source data and figure supplement(s) are available for figure 5: **Figure 5 - Source Data 1:** Ramos HCF-1_N_ annotated ChIP-Seq peaks. **Figure 5 - Source Data 2:** Annotated intersect of ChIP-Seq peaks for HCF-1_N_ and MYC in Ramos cells. **Figure 5 - figure supplement 1:** Comparison of MYC and HCF-1_N_ localization on Ramos cell chromatin. **Figure 5 - figure supplement 2:** HBM mutations do not impact DNA-binding by MYC:MAX complexes.

We previously reported that WDR5 has an important role in recruiting MYC to chromatin at a cohort of genes overtly linked to protein synthesis, including more than half of the ribosomal protein genes (Thomas et al., 2019). To determine whether these genes are also bound by HCF-1, we compared our HCF-1_N_ and MYC ChIP-Seq data to those we generated for WDR5 in this setting. Interestingly, there is little overlap of binding sites for MYC, WDR5, and HCF-1_N_ in Ramos cells, with just ~5% of MYC-HCF-1_N_ co-bound sites also being bound by WDR5 (Figure 5 - figure supplement 1H). Moreover, of the 88 sites bound by all three proteins, only three of these are sites where WDR5 has a functional role in MYC recruitment. Thus, despite the fact that WDR5 and HCF-1 are often both members of the same protein complex (Cai et al., 2010), and despite them both having links to key aspects of protein synthesis gene expression, the two proteins associate with MYC at distinct and separate regions of the genome.

Next, we overlaid the physical location of MYC and HCF-1_N_ on chromatin with gene expression changes we had monitored in earlier experiments. Looking at genes displaying promoter proximal binding of MYC and HCF-1_N_—where clear gene assignments can be made—we see that approximately one-third are differentially regulated in the presence of either the 4A or VP16 HBM MYC mutants (Figure 5G). For the 4A mutant, a slightly greater proportion of co-bound genes are down-regulated, while the opposite is apparent for the VP16 HBM mutant. A relatively small cohort of MYC–HCF-1_N_ co-bound genes are oppositely impacted by the 4A and VP16 HBM mutants (Figure 5G; “4A/VP”), but comparing these with those genes that are deregulated with depletion of HCF-1_N_ (Figure 5H), we again see that a majority are connected to ribosome biogenesis and mitochondria, and that most of these are positively regulated by HCF-1 and the MYC–HCF-1 interaction.

Finally, we asked if recruitment of MYC or HCF-1 to chromatin at a subset of genes from Figure 5H is altered by the 4A or VP16 HBM mutations. It has been reported that deletion of MbIV from N-MYC reduces the ability of MYC:MAX dimers to bind to DNA (Cowling et al., 2006). This phenotype is unrelated to the MYC HBM, however, as we determined that neither the 4A nor the VP16 HBM mutations have an overt impact on the binding of recombinant MYC:MAX dimers to DNA *in vitro* (Figure 5 - figure supplement 2A and 2B). Consistent with this result, we observe that the binding of MYC to chromatin in cells is not significantly affected by these mutations (Figure 5I). We also observe that binding of HCF-1 to these same sites is insensitive to HBM mutations in MYC (Figure 5J). The observation that binding of neither protein to chromatin is HBM-sensitive strongly supports the idea MYC and HCF-1 interact to control the expression of these genes through a co-recruitment-independent mechanism.

From these results, we conclude that, in this context, (i) MYC is a common binding partner with HCF-1 on chromatin, (ii) HCF-1 and WDR5 bind to different groups of MYC-bound genes, (iii) MYC and HCF-1 interact to directly promote the expression of genes connected to ribosome biogenesis and mitochondrial vigor, and (iv) gene expression changes that result directly from the MYC–HCF-1 interaction occur through a mechanism that is independent of their recruitment to chromatin.

### The MYC–HCF-1 interaction is important for tumor engraftment and maintenance

The ability of MYC to regulate ribosome (van Riggelen et al., 2010) and mitochondrial (Morrish and Hockenbery, 2014) biogenesis are core aspects of its tumorigenic repertoire. We would expect, therefore, that disrupting the MYC–HCF-1 interaction would have a significant impact on the ability of Ramos lymphoma cells to establish and maintain tumors *in vivo*. To address this expectation, we tested the impact of the 4A MYC mutant on tumorigenesis in mice. Because this is such an aggressive tumor model (Thomas et al., 2019), we did not test the gain-of-function VP16 HBM mutant. In these experiments, we included a second, independent, clone carrying the switchable 4A mutation (4A-1 and 4A-2); we also included a switchable Δ264 mutant (Thomas et al., 2019), which deletes residues in the carboxy-terminal half of MYC required for its nuclear localization, as well as interaction with WDR5, HCF-1, and MAX.

First, we assayed tumor engraftment by switching the engineered cells in culture and then injecting into the flanks of nude mice (Figure 6A). As expected, the WT to WT switched cells develop tumors rapidly *in vivo* (Figure 6B and Figure 6 - figure supplement 1A), resulting in all mice reaching humane endpoints and being euthanized by 21 days post-injection (Figure 6C). In contrast, 4A-1, 4A-2, and Δ264 switched cells are significantly delayed, both in tumor growth (Figure 6B and Figure 6 - figure supplement 1A) and mortality (Figure 6C). Although 4A-1, 4A-2, and Δ264 switched cells did form tumors, these appear to originate from the outgrowth of unswitched cells in the injected populations, as ~75% of cells in these tumors are in their unswitched state (Figure 6D). In this assay, therefore, the MYC–HCF-1 interaction is required for tumor growth, and there is little if any difference between disruption of the MYC–HCF-1 interaction and disabling the majority of the nuclear functions of MYC.

**Figure 6:**
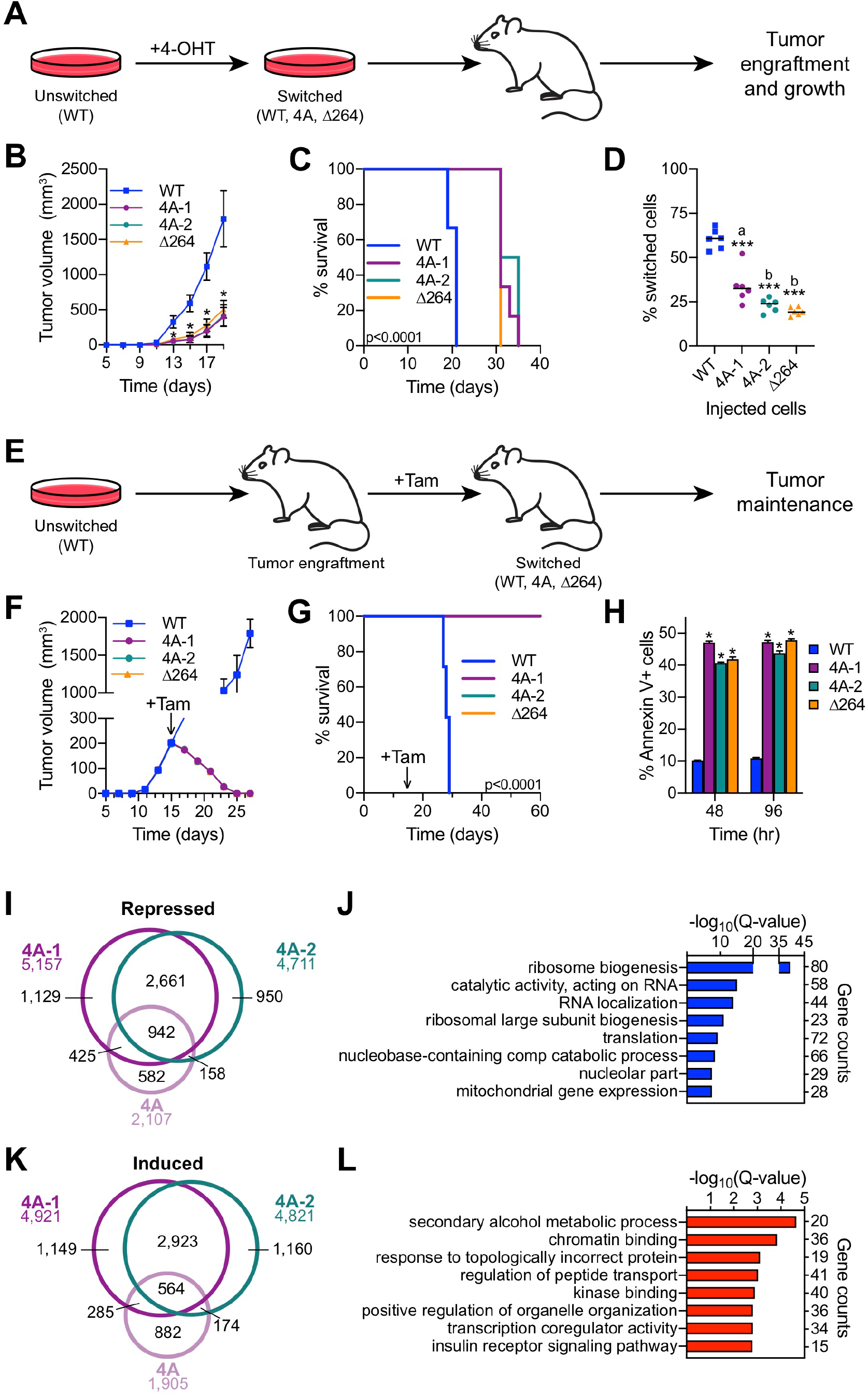
The MYC–HCF-1 interaction is required for tumor engraftment and maintenance. (**A**) Tumor engraftment schema: WT, 4A-1, 4A-2, and Δ264 cells are switched in culture prior to injection into a flank of nude mice to test the impact of the mutations on tumor engraftment and growth. (**B**) Average tumor volume over time following injection of switched cells. Shown are the mean and standard error for six mice each. Only days five to 19 are shown here; the full course of the experiment is depicted in **Figure 6 - figure supplement 2A**. Student’s t-test between WT and each of the mutants was used to calculate P-value; *P < 0.000043. (**C**) Kaplan-Meier survival curves of mice (n=6 of each) injected with switched cells. Log-rank test was used to calculate P-value (< 0.0001) from six biological replicates. (**D**) PCR assays of genomic DNA was used to determine the proportion of switched cells present in each tumor after sacrifice. Each dot represents an individual tumor, and the line indicates the mean for each group. Student’s t-test between WT and each of the mutants was used to calculate P-values; a = 0.0002, b < 0.0001. (**E**) Tumor maintenance schema: Unswitched WT, 4A-1, 4A-2, and Δ264 cells were injected into the flanks of nude mice. Tumors were grown until day 15, at which point mice received tamoxifen injections (one/day for three days) to induce switching of the cells. (**F**) Average tumor volume before and after cells were switched. The day at which tamoxifen (Tam) administration was initiated is indicated with an arrow. Shown are the mean and standard error for seven mice for WT and six mice for 4A-1, 4A-2, and Δ264 cells. (**G**) Kaplan-Meier survival curves of mice in the tumor maintenance assay (n=7 for WT, and n=6 for 4A-1, 4A-2, and Δ264). The day at which tamoxifen (Tam) administration was initiated is indicated with an arrow. Log-rank test was used to calculate P-value (< 0.0001). (**H**) Annexin V staining and flow cytometry were performed on cells isolated from tumors at 48 and 96 hours following the first tamoxifen administration to determine the extent of apoptosis. Shown are the mean and standard error for four mice each. Student’s t-test between WT and each of the mutants was used to calculate P-value; *P < 0.000001. (**I**) Venn diagram showing the relationship between genes significantly (FDR < 0.05) decreased in the 4A cell line and the 4A-1 and 4A-2 tumors. (**J**) Gene ontology enrichment analysis of genes significantly (FDR < 0.05) decreased in the 4A cell line, and the 4A-1 and 4A-2 tumors. (**K**) Venn diagram showing the overlap of genes significantly (FDR < 0.05) increased in the 4A cell line, and the 4A-1 and 4A-2 tumors. (**L**) Gene ontology enrichment analysis of genes significantly (FDR < 0.05) increased in the 4A cell line, and the 4A-1 and 4A-2 tumors. The following source data and figure supplement(s) are available for figure 6: **Figure 6 - Source Data 1:** Tumor RNA-Seq significant changes. **Figure 6 - figure supplement 1:** Disruption of the MYC−HCF-1 interaction *in vivo*.

Next, we injected unswitched cells into the flanks of mice, allowed tumors to form, and then switched to each of the MYC variants by injecting mice with tamoxifen (Figure 6E). As we observed previously (Thomas et al., 2019), the “WT to WT” tumors continue to grow rapidly after switching (Figure 6F and Figure 6 - figure supplement 1B), and all mice had to be euthanized before 30 days (Figure 6G). For the 4A switches, however, tumors rapidly regressed (Figure 6F), and all mice survived—and were tumor free—for the 60 day duration of the experiment (Figure 6 - figure supplement 1B). Regression of the 4A tumors occurs at a pace that is virtually indistinguishable from the Δ264 mutant (Figure 6 - figure supplement 1B), and like the Δ264 scenario, is accompanied by high levels of apoptosis, as measured by Annexin V staining (Figure 6H), caspase activity (Figure 6 - figure supplement 1C), and sub-G_1_ DNA content (Figure 6 - figure supplement 1D). We conclude that the interaction of MYC with HCF-1 is essential for tumor maintenance in this context.

Finally, we performed RNA-Seq on tumor material excised 48 hours after switching (Figure 6 - figure supplement 1E, Figure 6 - Source Data 1). Thousands of differentially expressed genes were identified in the mutant switch tumors, many of which are shared between the 4A and Δ264 mutants—for both the repressed (Figure 6 - figure supplement 1F) and induced (Figure 6 - figure supplement 1G) directions. Common reduced transcripts are enriched in those connected to ribosome biogenesis, translation, and mitochondrial envelope (Figure 6 - figure supplement 1H), whereas those that are induced are enriched for functions including transcription co-regulator activity, kinase binding, and the vacuole (Figure 6 - figure supplement 1I). Many of these gene expression changes are likely due to indirect effects of tumor regression, and so to focus on those connected to the MYC–HCF-1 interaction, we overlaid tumor RNA-Seq with that generated for the 4A MYC mutant *in vitro* (Figure 3). Interestingly, more than 70% of the genes repressed in the 4A cell line are also repressed in either 4A-1 or 4A-2 tumors, and there is a common set of 942 genes that are shared between all three datasets (Figure 6I). These genes coalesce on those connected to ribosome biogenesis, translation, and the mitochondria (Figure 6J). The overlap of induced genes was less pronounced (30%; Figure 6K) and these genes are less clustered, although we do observe modest enrichment in categories connected to metabolism, chromatin binding, and transcription coregulator activity (Figure 6L). The recurring connections we observe between the MYC–HCF-1 interaction and ribosome biogenesis and mitochondrial function, both *in vitro* and *in vivo*, strongly supports the notion that a major function of this interaction is stimulate ribosome production and mitochondrial vigor, and that these actions are central for the ability of MYC to drive tumor onset and maintenance.

## DISCUSSION

The wealth of MYC-interaction partners provides a rich resource for the discovery of novel ways to eventually inhibit MYC in the clinic. Unfortunately, the complexity of the MYC interactome also presents a barrier to prioritizing which co-factors to pursue. The highest priority co-factors should be those that directly interact with MYC, play a critical role in the core tumorigenic functions of the protein, and where there is proof-of-concept that disrupting interaction with MYC would provide a therapeutic benefit in the context of an existing malignancy. Here, we provide this information for HCF-1. We show that MYC directly interacts with HCF-1 via a non-canonical HBM, identify roles for the MYC–HCF-1 interaction in the control of genes involved in ribosome biogenesis, translation, and mitochondrial function, define the impact of modulation of this interaction on the metabolome, and show that loss of the MYC–HCF-1 interaction promotes frank and irreversible tumor regression *in vivo*. Although we do not yet know if a therapeutic window exists for targeting MYC through HCF-1, these findings cement HCF-1 as a critical MYC co-factor and one worth pursuing as means to inhibit MYC in cancer.

Through the use of mutations that bidirectionally modulate the interaction between MYC and HCF-1, inducible degradation of HCF-1_N_, and ChIP-Seq analyses, we identified a relatively small set of genes that we posit are direct targets of the MYC–HCF-1 interaction. Given the stringency of our approach, the list of genes we derive is likely an underestimate of the totality of direct MYC–HCF-1 targets. But what is particularly interesting about this set are their biological clustering, their connections to core pro-tumorigenic MYC activities, and their ability to account for many of the overt consequences of modulating the MYC–HCF-1 interaction on metabolism and tumorigenesis. What is also interesting about this set of genes is that they appear to be regulated by MYC and HCF-1 through a co-recruitment-independent mechanism.

Among the direct targets of the MYC−HCF-1 interaction are genes that catalyze rate-limiting steps in both ribosome biogenesis (rDNA transcription by POLR1A) and translation (initiator tRNA binding to start codon by EIF2S1 [EIF-2α]) (Hershey, 1991; Laferte et al., 2006). MYC regulates ribosome biogenesis through controlling and coordinating the transcription of ribosomal DNA, ribosomal protein genes, and components in the processing and assembly of ribosomes (van Riggelen et al., 2010). Interestingly, even with the direct MYC–HCF-targets classified as mitochondrially-connected, we see links to protein synthesis—MARS2, MRPS12, and MRPL32, for example, are specifically involved in the synthesis of mitochondrial proteins, including those required for oxidative phosphorylation. We conclude that, in the context of its relationship with MYC, HCF-1 is dedicated to promoting multiple aspects of biomass accumulation.

The concept that there can be process-specific co-factors for MYC is not widely appreciated. Attention is generally placed on those co-factors required for general functioning of the MYC protein. But our studies of HCF-1—as well as our earlier work on WDR5 (Thomas et al., 2019)—establish the precedent that such dedicated co-factors do exist. Indeed, there are interesting parallels between the interaction of MYC with HCF-1 and with WDR5—both are mediated by conserved MYC boxes, both control the expression of relatively small cohorts of genes, both are required to establish and maintain tumors, and both have clear links to biomass accumulation. Beyond this point, however, these parallels break down. There is little overlap of target genes regulated by the MYC–WDR5 and MYC–HCF-1 interactions, and whereas MYC and HCF-1 associate to predominantly control ribosome biogenesis, MYC and WDR5 work together to stimulate the expression of genes encoding structural components of the ribosome (Thomas et al., 2019). This division of labor between HCF-1 and WDR5 in different aspects of protein synthesis is particularly intriguing given that HCF-1 and WDR5 work together as part of multi-protein chromatin modifying complexes (Tyagi et al., 2007; Wysocka et al., 2003). Yet MYC interacts with each protein through separate MYC boxes, and indeed the way in which MYC interacts with WDR5 likely precludes WDR5 from assembling into canonical HCF-1-containing complexes (Thomas et al., 2015). We believe the separation of interaction surfaces allows MYC to access non-canonical functions of both WDR5 and HCF-1, and may have evolved to permit discrete regulation of the constituents of the ribosomes versus the factors required to assemble these constituents into ribosome particles. Indeed, because HCF-1 (Mazars et al., 2010), MYC (Chou et al., 1995), and the MYC–HCF-1 interaction (Itkonen et al., 2019) all have ties to the metabolic sensor OGT (Swamy et al., 2016), it is possible that this separation of function allows for rapid modulation of ribosome assembly by HCF-1 when the metabolic state of the cell declines, while at the same time ensuring that ribosome subunits are present for rapid recovery when metabolism ramps up.

The concept that HCF-1 is a biomass-specific co-factor for MYC can also account for our discovery that intracellular amino acid levels are increased when the MYC–HCF-1 interaction is disrupted. Impairment of ribosome biogenesis and translation can lead to an accumulation of amino acids and compensatory changes in the expression of their transporters (Guan et al., 2014; Scott et al., 2014). Of note, the way in which amino acid levels almost globally respond to changes in the MYC–HCF-1 interaction can also provide a simple explanation for how the MYC–HCF-1 interaction contributes to the glutamine addiction of Ramos cells. If MYC partners with HCF-1 to drive biomass accumulation, the net effect of this interaction will be to increase the demand for intracellular amino acids, with glutamine playing a particularly important role in the biosynthesis of multiple non-essential amino acids, including glutamate, arginine, proline, and alanine (Hosios et al., 2016). The ability of MYC to drive glutamine-addiction is a defining characteristic of its tumorigenic repertoire (Tansey, 2014), and is thought to result from the ability of MYC to promote glutaminolysis and induce the expression of amino acid transporters (Wise et al., 2008). But the accumulation of glutamine we observe upon disruption of the MYC–HCF-1 interaction, together with the specific role HCF-1 plays in MYC function, suggests that glutamine addiction might also be fueled by the well-established ability of MYC to stimulate protein synthesis (Iritani and Eisenman, 1999).

Our discovery that MYC and HCF-1 work together to regulate transcription via a co-recruitment-independent mechanism is surprising, but not without precedent. E2F transcription factors interact with HCF-1 through a canonical HBM to control the expression of genes connected to cell cycle progression, yet depletion of these proteins has little effect on the recruitment of either to chromatin (Iwata et al., 2013; Parker et al., 2014; Tyagi et al., 2007). Similarly, *Drosophila* Myc and HCF interact to activate transcription and control growth, but their respective interactions with chromatin are independent of one another (Furrer et al., 2010). A recruitment-independent mechanism may thus be a common way in which transcription factors interact with HCF-1 to modulate transcription.

Finally, our demonstration that disrupting the MYC–HCF-1 interaction in the context of an existing tumor promotes its regression provides compelling proof-of-concept for the idea that inhibitors of this interaction could have utility as anti-cancer agents. Switching WT MYC to the 4A mutant caused rapid and widespread induction of apoptosis, and was associated with changes in the expression of genes connected to ribosome biogenesis, translation, and the mitochondria, consistent with the idea that reduced expression of these MYC–HCF-1 target genes triggers the regression process. The small and well-defined nature of the HBM suggests that, if structural information becomes available for the HCF-1 VIC domain, it could be possible to develop small molecule inhibitors that block the MYC–HCF-1 interaction. The most obvious concern with this strategy is that HCF-1 is not a MYC-specific co-factor, and that its interactions with other transcription factors may limit or prevent attainment of a therapeutic window. To our knowledge, MYC proteins and E2F3a are the only transcription factors that interact with HCF-1 via an “imperfect” HBM, which we show here is sub-optimal for robust HCF-1 association. It might be possible to develop a therapeutic window by exploiting the non-canonical nature of the HBM in MYC, with the expectation that this interaction will be more sensitive than others that carry higher affinity HBM motifs. We note, however, that many of the factors with which HCF-1 interacts via an HBM are inherently pro-proliferative, with the E2F proteins in particular playing a predominant role in cancer initiation, maintenance, and response to therapies (Kent and Leone, 2019). We also note that HCF-1 has been reported to be overexpressed in cancer, and its overexpression can correlate with poor clinical outcomes (Glinsky et al., 2005). It is possible, therefore, that on-target collateral consequences of inhibiting the MYC–HCF-1 interaction could also have therapeutic benefit against malignancies.

## MATERIALS AND METHODS

### Key Resources table

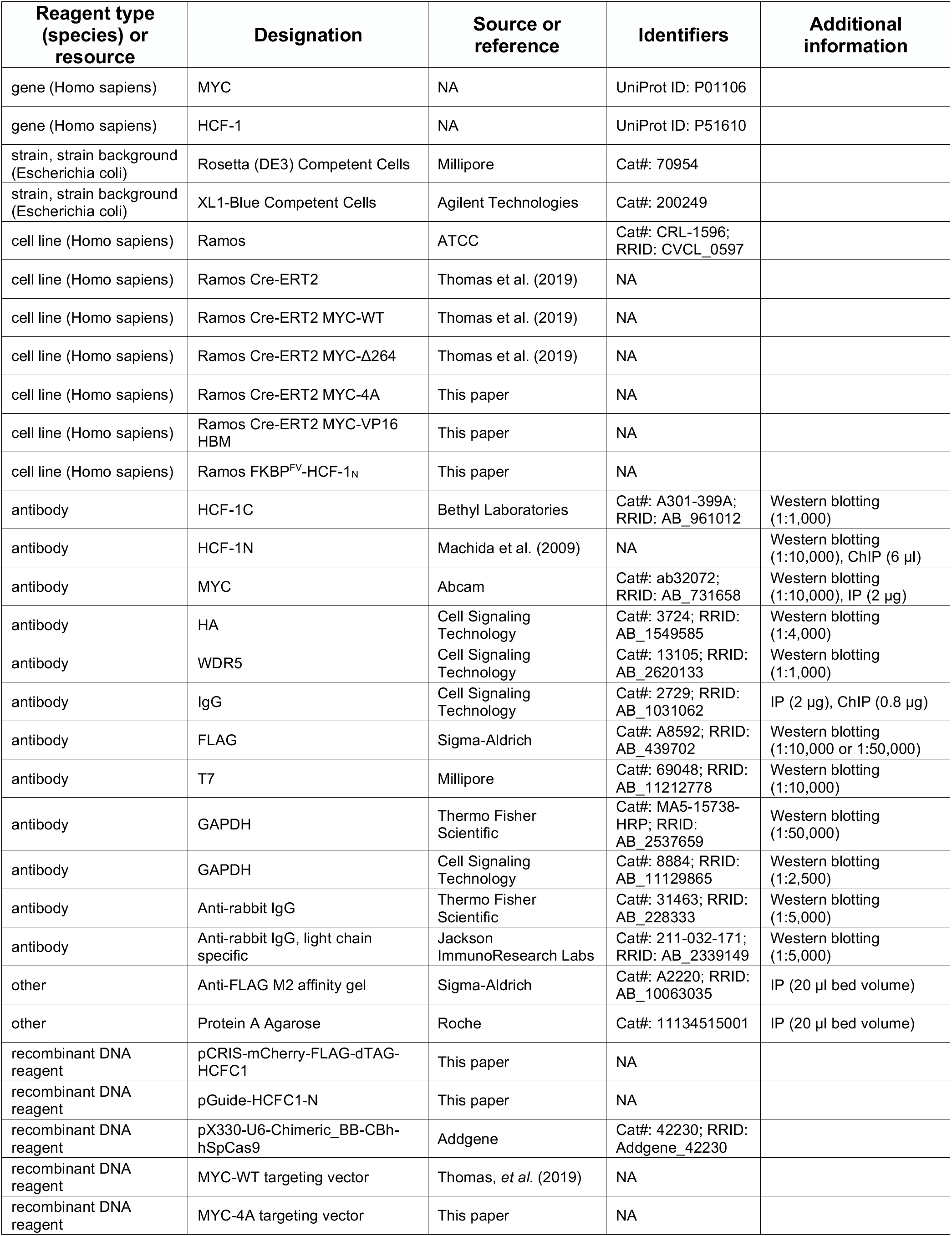
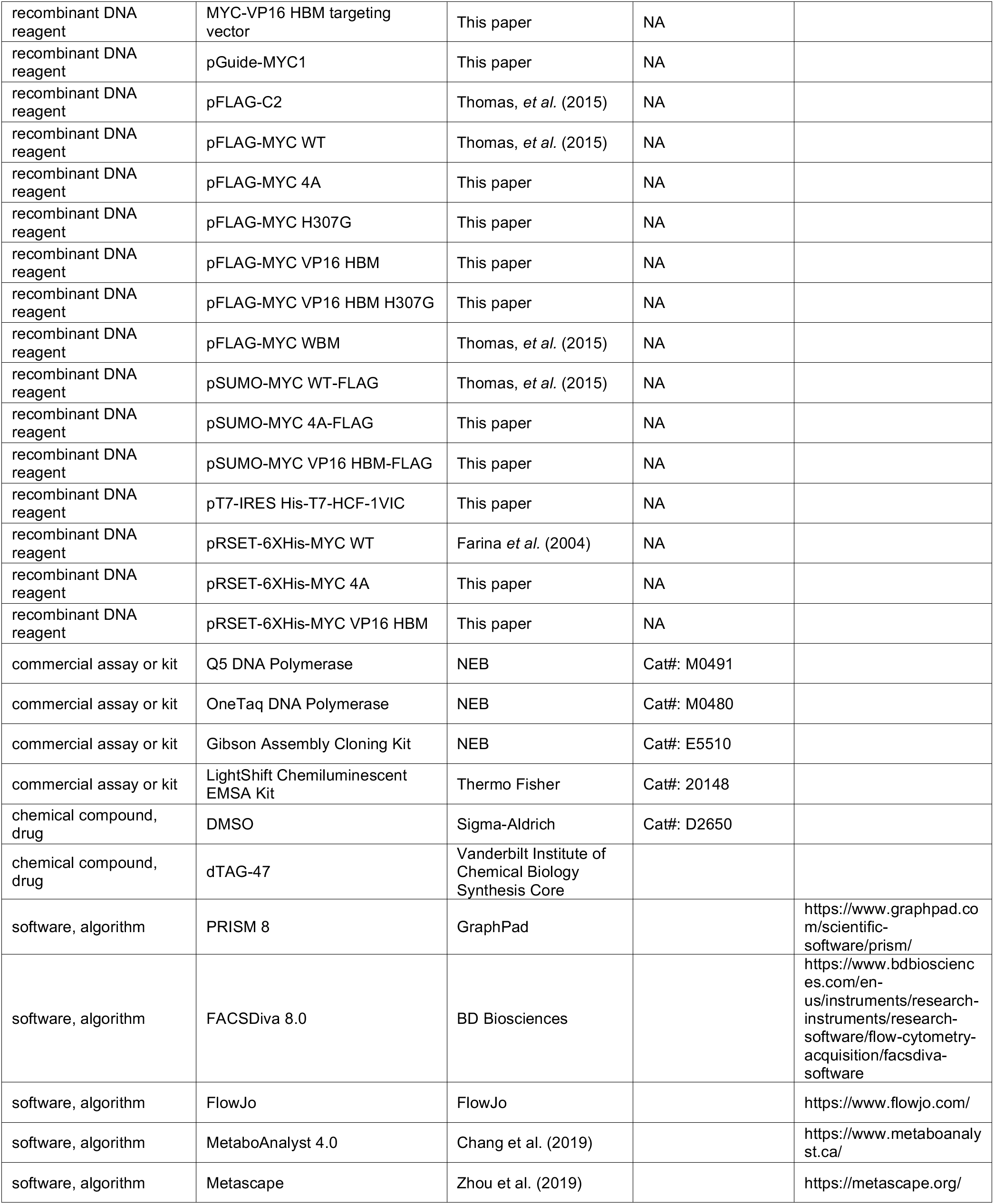

| Reagent type (species) or resource | Designation | Source or reference | Identifiers | Additional information |
| --- | --- | --- | --- | --- |
| gene (Homo sapiens) | MYC | NA | UniProt ID: P01106 |  |
| gene (Homo sapiens) | HCF-1 | NA | UniProt ID: P51610 |  |
| strain, strain background (Escherichia coli) | Rosetta (DE3) Competent Cells | Millipore | Cat#: 70954 |  |
| strain, strain background (Escherichia coli) | XL1-Blue Competent Cells | Agilent Technologies | Cat#: 200249 |  |
| cell line (Homo sapiens) | Ramos | ATCC | Cat#: CRL-1596; RRID: CVCL_0597 |  |
| cell line (Homo sapiens) | Ramos Cre-ERT2 | Thomas et al. (2019) | NA |  |
| cell line (Homo sapiens) | Ramos Cre-ERT2 MYC-WT | Thomas et al. (2019) | NA |  |
| cell line (Homo sapiens) | Ramos Cre-ERT2 MYC-Δ264 | Thomas et al. (2019) | NA |  |
| cell line (Homo sapiens) | Ramos Cre-ERT2 MYC-4A | This paper | NA |  |
| cell line (Homo sapiens) | Ramos Cre-ERT2 MYC-VP16 HBM | This paper | NA |  |
| cell line (Homo sapiens) | Ramos FKBP<sup>FV</sup>-HCF-1<sub>N</sub> | This paper | NA |  |
| antibody | HCF-1C | Bethyl Laboratories | Cat#: A301-399A; RRID: AB_961012 | Western blotting (1:1,000) |
| antibody | HCF-1N | Machida et al. (2009) | NA | Western blotting (1:10,000), ChIP (6 μl) |
| antibody | MYC | Abcam | Cat#: ab32072; RRID: AB_731658 | Western blotting (1:10,000), IP (2 μg) |
| antibody | HA | Cell Signaling Technology | Cat#: 3724; RRID: AB_1549585 | Western blotting (1:4,000) |
| antibody | WDR5 | Cell Signaling Technology | Cat#: 13105; RRID: AB_2620133 | Western blotting (1:1,000) |
| antibody | IgG | Cell Signaling Technology | Cat#: 2729; RRID: AB_1031062 | IP (2 μg), ChIP (0.8 μg) |
| antibody | FLAG | Sigma-Aldrich | Cat#: A8592; RRID: AB_439702 | Western blotting (1:10,000 or 1:50,000) |
| antibody | T7 | Millipore | Cat#: 69048; RRID: AB_11212778 | Western blotting (1:10,000) |
| antibody | GAPDH | Thermo Fisher Scientific | Cat#: MA5-15738-HRP; RRID: AB_2537659 | Western blotting (1:50,000) |
| antibody | GAPDH | Cell Signaling Technology | Cat#: 8884; RRID: AB_11129865 | Western blotting (1:2,500) |
| antibody | Anti-rabbit IgG | Thermo Fisher Scientific | Cat#: 31463; RRID: AB_228333 | Western blotting (1:5,000) |
| antibody | Anti-rabbit IgG, light chain specific | Jackson ImmunoResearch Labs | Cat#: 211-032-171; RRID: AB_2339149 | Western blotting (1:5,000) |
| other | Anti-FLAG M2 affinity gel | Sigma-Aldrich | Cat#: A2220; RRID: AB_10063035 | IP (20 μl bed volume) |
| other | Protein A Agarose | Roche | Cat#: 11134515001 | IP (20 μl bed volume) |
| recombinant DNA reagent | pCRIS-mCherry-FLAG-dTAG-HCFC1 | This paper | NA |  |
| recombinant DNA reagent | pGuide-HCFC1-N | This paper | NA |  |
| recombinant DNA reagent | pX330-U6-Chimeric_BB-CBh-hSpCas9 | Addgene | Cat#: 42230; RRID: Addgene_42230 |  |
| recombinant DNA reagent | MYC-WT targeting vector | Thomas, *et al.* (2019) | NA |  |
| recombinant DNA reagent | MYC-4A targeting vector | This paper | NA |  |
| recombinant DNA reagent | MYC-VP16 HBM targeting vector | This paper | NA |  |
| recombinant DNA reagent | pGuide-MYC1 | This paper | NA |  |
| recombinant DNA reagent | pFLAG-C2 | Thomas, *et al.* (2015) | NA |  |
| recombinant DNA reagent | pFLAG-MYC WT | Thomas, *et al.* (2015) | NA |  |
| recombinant DNA reagent | pFLAG-MYC 4A | This paper | NA |  |
| recombinant DNA reagent | pFLAG-MYC H307G | This paper | NA |  |
| recombinant DNA reagent | pFLAG-MYC VP16 HBM | This paper | NA |  |
| recombinant DNA reagent | pFLAG-MYC VP16 HBM H307G | This paper | NA |  |
| recombinant DNA reagent | pFLAG-MYC WBM | Thomas, *et al.* (2015) | NA |  |
| recombinant DNA reagent | pSUMO-MYC WT-FLAG | Thomas, *et al.* (2015) | NA |  |
| recombinant DNA reagent | pSUMO-MYC 4A-FLAG | This paper | NA |  |
| recombinant DNA reagent | pSUMO-MYC VP16 HBM-FLAG | This paper | NA |  |
| recombinant DNA reagent | pT7-IRES His-T7-HCF-1VIC | This paper | NA |  |
| recombinant DNA reagent | pRSET-6XHis-MYC WT | Farina *et al.* (2004) | NA |  |
| recombinant DNA reagent | pRSET-6XHis-MYC 4A | This paper | NA |  |
| recombinant DNA reagent | pRSET-6XHis-MYC VP16 HBM | This paper | NA |  |
| commercial assay or kit | Q5 DNA Polymerase | NEB | Cat#: M0491 |  |
| commercial assay or kit | OneTaq DNA Polymerase | NEB | Cat#: M0480 |  |
| commercial assay or kit | Gibson Assembly Cloning Kit | NEB | Cat#: E5510 |  |
| commercial assay or kit | LightShift Chemiluminescent EMSA Kit | Thermo Fisher | Cat#: 20148 |  |
| chemical compound, drug | DMSO | Sigma-Aldrich | Cat#: D2650 |  |
| chemical compound, drug | dTAG-47 | Vanderbilt Institute of Chemical Biology Synthesis Core |  |  |
| software, algorithm | PRISM 8 | GraphPad |  | https://www.graphpad.com/scientific-software/prism/ |
| software, algorithm | FACSDiva 8.0 | BD Biosciences |  | https://www.bdbiosciences.com/en-us/instruments/research-instruments/research-software/flow-cytometry-acquisition/facsdiva-software |
| software, algorithm | FlowJo | FlowJo |  | https://www.flowjo.com/ |
| software, algorithm | MetaboAnalyst 4.0 | Chang et al. (2019) |  | https://www.metaboanalyst.ca/ |
| software, algorithm | Metascape | Zhou et al. (2019) |  | https://metascape.org/ |

### Primers and cloning

See Supplementary File 1 for primer sequences. PCRs were performed using either Q5 DNA Polymerase (NEB, Ipswich, Massachusetts, M0491) or OneTaq DNA Polymerase (NEB M0480). Gibson assemblies were performed using Gibson Assembly Cloning Kit (NEB E5510). More specific details about cloning steps can be found in the relevant sections.

### Cell culture and transient transfections

293T cells were maintained in DMEM with 4.5 g/l glucose, L-Glutamine, and sodium pyruvate (Corning, Corning, New York, 10-013-CV), and supplemented with 10% fetal bovine serum (FBS, Denville Scientific, Metuchen, New Jersey, FB5001-H), and 1% penicillin/streptomycin (P/S, Gibco, Waltham, Massachusetts, 15140122). Ramos cells were obtained from the ATCC (Manassas, Virginia, CRL-1596) and maintained in RPMI 1640 with L-glutamine (Corning 10-040-CV), and supplemented with 10% FBS (Denville Scientific FB5001-H), and 1% P/S (Gibco 15140122).

### Antibodies

Rabbit anti-HCF1_C_ polyclonal (Bethyl Laboratories, Montgomery, Texas, Cat# A301-399A); rabbit anti-HCF1_N_ polyclonal (Machida et al., 2009); rabbit anti-c-MYC monoclonal (Abcam, Cambridge, United Kingdom, Cat# ab32072); rabbit anti-HA (C29F4) monoclonal (Cell Signaling Technology, Danvers, Massachusetts, Cat# 3724); rabbit anti-WDR5 (D9E1I) monoclonal (Cell Signaling Technology Cat# 13105); rabbit anti-IgG polyclonal (Cell Signaling Technology Cat# 2729); mouse anti-FLAG monoclonal, HRP-conjugated (Sigma-Aldrich, St. Louis, Missouri, Cat# A8592); mouse anti-T7 monoclonal, HRP conjugated (Millipore, Burlington, Massachusetts, Cat# 69048); mouse anti-GAPDH monoclonal, HRP conjugated (Thermo Fisher Scientific, Waltham, Massachusetts, Cat# MA5-15738-HRP); rabbit anti-GAPDH (D16H11) monoclonal, HRP conjugated (Cell Signaling Technology Cat# 8884); goat anti-rabbit IgG monoclonal, HRP conjugated (Thermo Fisher Scientific Cat# 31463); mouse anti-rabbit IgG monoclonal, light chain specific, HRP conjugated (Jackson ImmunoResearch Laboratories, West Grove, Pennsylvania, Cat# 211-032-171).

### Generation of switchable MYC allele Ramos cell lines

Q5 site-directed mutagenesis of MYC-WT targeting vector from Thomas *et al.* (2019) was used to create MYC-4A (4A_F and 4A_R) and MYC-VP16 HBM (VP16 HBM_F and VP16 HBM_R). The pGuide plasmid described by Thomas *et al.* (2019) was used as a backbone to introduce the sgRNA sequence GCTACGGAACTCTTGTGCGTA (pGuide-MYC1) by Q5 site directed mutagenesis with the primers GUIDE MYC-1A and GUIDE MYC-1B.

For the generation of switchable cells, 10 million Ramos cells stably expressing CRE-ER^T2^ (Thomas et al., 2019) were electroporated (BioRad Gene Pulser II, 220 V and 950 μF) with 10 μg of relevant targeting vector (MYC-4A or MYC-VP16 HBM), 15 μg pGuide-MYC1, and 15μ g pX330-U6-Chimeric_BB-CBh-hSpCas9 (gift from Feng Zhang, AddGene plasmid #42230) (Cong et al., 2013). WT and Δ264 cell lines were the same as those used in Thomas *et al.* (2019). Cells were treated with 150 ng/ml puromycin (Sigma-Aldrich P7255) and 100 μg/ml hygromycin (Corning 30240CR), selecting for the switchable MYC cassette and CRE-ER^T2^ recombinase, respectively. Following selection, single cells, stained using propidium iodide (PI, Sigma-Aldrich P4864) for viability, were sorted by the Vanderbilt Flow Cytometry Shared Resource using a BD FACSAria III flow cytometer into a 96-well plate to generate clonal cell lines under puromycin and hygromycin selection. Individual clones were expanded, and initially validated by switching for 24 hours using 20 nM (Z)-4-Hydroxytamoxifen (4-OHT, Tocris, Minneapolis, Minnesota, 3412), and flow cytometry (see below) for GFP expression. Further validation was performed by western blotting after 24 hours 20 nM ±4-OHT (see below), and by Southern blotting (see below). The 4A cell line (4A-1) used for the majority of experiments is haploinsufficient for part of chromosome 11, approximately between co-ordinates 118,685,194 and 134,982,408. This region of the genome was excluded from genomic analyses. For all experiments, switching was performed by treatment with 20 nM 4-OHT for 2 or 24 hours (see relevant method or figure legend).

### Generation of dTAG Ramos cell lines

To create pCRIS-mCherry-FLAG-dTAG-HCFC1, pCRIS-PITChv2-Puro-dTAG (BRD4) (gift from James Bradner, AddGene plasmid #91793) (Nabet et al., 2018) was first modified to remove the BRD4 homology arms and replace the 2XHA tags with a FLAG tag. This was done by Gibson assembly of the vector (Q5 amplification using pCRIS-HCFC1N_F and pCRIS_R), puromycin cassette (Q5 amplification using Puro-1 and Puro-FLAG_R), and FKBP12^FV^ (Q5 amplification using FLAG-FKBP_F and pCRIS-FKBP_R). The resulting vector was again modified using Gibson assembly by combining the vector (Q5 amplification using pCRIS-HCFC1N_F and pCRIS-HCFC1N_R), an upstream 271bp *HCFC1* 5 ‘homology arm (hg19 chrX:153236265-153236535, OneTaq amplification using HCFC1N_F and HCFC1N-5’Hom_R), mCherry (Q5 amplification using mCherry_F and mCherry_R), FKBP12^FV^ (Q5 amplification using HCFC1-mCherry-FKBP_F and HCFC1-mCherry-FKBP_R), and a downstream 800bp *HCFC1* 3’ homology arm (hg19 chrX:153235465-153236264, OneTaq amplification using HCFC1N-3’Hom_F and HCFC1N-3’Hom_R). The pGuide plasmid described by Thomas *et al.* (2019) was used as a backbone to introduce the guide RNA sequence CAGAAGCACCGCTGGCAAGT (pGuide-HCFC1-N) by Q5 site-directed mutagenesis with the primers HCFC1N-sgRNA_F and HCFC1N-sgRNA_R.

Fifteen micrograms pGuide-HCFC1-N and 10 μg pCRIS-mCherry-FLAG-dTAG-HCFC1 were electroporated (BioRad, Hercules, California, Gene Pulser II, 220 V and 950 μF) into Ramos cells, alongside 15 μg pX330-U6-Chimeric_BB-CBh-hSpCas9 (gift from Feng Zhang, AddGene plasmid #42230) (Cong et al., 2013) into 10 million Ramos cells. Because *HCFC1* is on the X chromosome and Ramos cells are XY (Klein et al., 1975), only a single copy of *HCFC1* is present for targeting using CRISPR/Cas9. Following electroporation, cells were expanded and a population of mCherry-positive cells, stained using Zombie NIR viability dye (BioLegend, San Diego, California, 423105), was sorted by the Vanderbilt Flow Cytometry Shared Resource using a FACSAria III flow cytometer (Becton Dickinson (BD), Franklin Lakes, New Jersey). This population of cells was expanded further before validation by western blotting. All experiments were conducted using this population, and were treated with either DMSO (Sigma-Aldrich D2650) or 500 nM dTAG-47, which was synthesized by the Vanderbilt Institute of Chemical Biology Synthesis Core.

### Transient transfection, western blotting and immunoprecipitation

Q5 site-directed mutagenesis of pFLAG-MYC WT (Thomas et al., 2015) was used to generate the following plasmids: pFLAG-MYC 4A (4A_F and 4A_R), pFLAG-MYC H307G (H307G_F and H307G_R), pFLAG-MYC VP16 HBM (VP16 HBM_F and VP16 HBM_R), and pFLAG-MYC VP16 HBM H307G (VP16 HBM H307G_F and VP16 HBM H307G_R). pFLAG-MYC WBM is from Thomas *et al.* (2015). Plasmid (19 μg) was prepared with 0.25 M CaCl_2_, incubated for 10 minutes with 1X HBS (2X HBS: 140 mM NaCl, 1.5 mM Na_2_HPO_4_, 50 mM HEPES, pH to 7.05), then applied drop-wise to 293T cells. Cells were grown for two days before harvesting (see below).

Cell lysates for western blotting or immunoprecipitation (IP) were prepared by rinsing cells twice in ice-cold 1X PBS, and harvesting in Kischkel Buffer (50 mM Tris pH 8.0, 150 mM NaCl, 5 mM EDTA, 1% Triton X-100) + protease inhibitor cocktail (PIC, Roche, Basel, Switzerland, 05056489001). Cells were sonicated at 25% power for 10 s (Cole-Parmer, Vernon Hills, Illinois, GE 130PB-1), debris removed by centrifugation, and protein concentration determined using Protein Assay Dye (BioRad 500-0006) against a bovine serum albumin (BSA) standard. For western blotting, lysate was diluted in 5X Laemmli Buffer (375 mM Tris pH 6.8, 40% glycerol, 10% SDS, bromophenol blue, 2-Mercaptoethanol). For IP, the concentration of samples was balanced using Kischkel Buffer. Antibody or anti-FLAG M2 Affinity Gel (Millipore A2220) was added, and samples rotated overnight at 4°C. For unconjugated antibodies, 20 μl bed volume of Protein A Agarose (Roche 11134515001) was added the following day to each sample, and rotated for 2-4 hours at 4°C. Samples were then washed 4X with Kischkel buffer (2X 4°C and 2X at room-temperature), and incubated in Laemmli buffer for 5 minutes at 95°C.

Protein from lysates and IPs were separated out by SDS-polyacrylamide gel electrophoresis (PAGE) in running buffer (25 mM Tris, 192 mM glycine, 0.2% SDS). Wet transfer to PVDF (PerkinElmer, Waltham, Massachusetts, NEF1002) was carried out in Towbin Transfer Buffer (25 mM Tris, 192 mM glycine, and 10% methanol). Membrane was blocked in 5% milk in TBS-T (20 mM Tris pH 7.6, 140 mM NaCl, 0.1% Tween-20), hybridized overnight in primary antibody (or 1 hour for HRP-conjugated), and for 1 hour in HRP-conjugated secondary antibody (if required). ECL substrates, SuperSignal West Pico (Pierce, Waltham, Massachusetts, 34080), Pico+ (Pierce 34580) and Femto (Pierce 34095), were used in various combinations for detection of bands by exposure to film.

### Flow cytometry and cell cycle analysis

Cells were filtered into 35 μm nylon mesh Falcon round bottom test tubes for flow cytometry, which was performed in the Vanderbilt Flow Cytometry Shared Resource. Single cells were gated based on side and forward scatter (for example, see Supplementary File 2) using the stated instruments.

To determine the proportion of switching of the switchable MYC allele Ramos cell lines, cells were fixed in 1% formaldehyde (FA, Thermo Fisher 28908) in PBS for 10 minutes at room temperature. The number of GFP-positive cells was determined using a BD LSR II flow cytometer.

For cell cycle analysis of the switchable MYC allele Ramos cell lines, cells were treated with 4-OHT for 2 hours, at which point the media was replaced. Cells were maintained for 7 days, at which point 1×10^6^ cells were collected and fixed in 1% FA (Thermo Fisher 28908) in PBS for 10 minutes at room temperature, then washed 2X with PBS. Permeabilization and staining was done using cell cycle staining buffer (PBS, 10 μg/ml PI (Sigma-Aldrich P4864), 100 μg/ml RNAse A, 2 mM MgCl_2_, 0.1% Triton X-100) for 25 minutes at room temperature, then stored overnight at 4°C. PI staining of at least 10,000 single cells for each the GFP-negative (GFP-) and GFP-positive (GFP+) populations was measured using a BD LSR Fortessa, and cell cycle distribution determined using BD FACSDIVA software

For cell cycle analysis of the FKBP^FV^-HCF-1_N_ Ramos cells, cells were treated with DMSO or 500 nM dTAG for 24 hours. One million cells were collected and fixed in ethanol overnight. After fixation, cells were stained overnight at 4°C in cell cycle staining buffer (PBS, 10 μg/ml PI (Sigma-Aldrich P4864), 100 μg/ml RNAse A, 2 mM MgCl2). PI staining of at least 10,000 single cells was measured using a BD LSR Fortessa, and cell cycle distribution determined using BD FACSDIVA software.

See section *In vivo* studies: Tumor formation and maintenance assays for details regarding flow cytometry experiments conducted on cells extracted from mice tumors.

### Southern blotting

Genomic DNA (gDNA) was prepared from parental and unswitched MYC switchable cells (WT, 4A, and VP16 HBM) (Miller et al., 1988). Briefly, cells were rinsed in ice-cold 1X PBS, and resuspended in DNA extraction buffer (10 mM Tris pH 8.1, 400 mM NaCl, 10 mM EDTA, 1% SDS, 50 μg/ml proteinase K (PK, Macherey-Nagel, Düren, Germany, 740506)). Lysis was performed overnight in a rotisserie at 56°C, before gDNA was extracted using ethanol precipitation. Southern blot was performed similarly to that described by Southern *et al*. (1975). gDNA (10 μg) was digested using XbaI (NEB R0145) and run out on a 1% agarose gel. DNA was transferred overnight to Hybond-N+ nylon membrane (GE Healthcare, Chicago, Illinois, RPN303B) by capillary action in transfer buffer (0.5 M NaOH, 0.6 M NaCl). The following day the membrane was immersed in neutralization buffer (1 M NaCl, 0.5 M Tris pH 7.4), UV cross-linked, and pre-hybridized overnight at 42°C in pre-hybridization buffer (50% formamide, 5X SSCPE (20X SSCPE: 2.4 M NaCl, 0.3 M Na citrate, 0.2 M KH2PO4, 0.02 M EDTA), 5X Denhardt’s solution (Invitrogen, Waltham, Massachusetts, 750018), 0.5 mg/ml salmon sperm DNA (Agilent, Santa Clara, California, 201190), 1% SDS). Templates for probe generation were prepared by Q5 amplification from MYC-WT targeting vector using primers GFP_F and GFP_R (GFP template) and from parental Ramos cell gDNA using primers 5’_F and 5’_R, followed by gel purification. Probes were prepared by random priming of corresponding PCR products (5’ and GFP templates) in the presence of [αP32]CTP (PerkinElmer BLU513H100UC). Unincorporated nucleotides were removed using a Sephadex G-50 column (GE Healthcare 28-9034-08). The membrane was incubated overnight at 42°C with probe in hybridization buffer (50% formamide, 5X SSCPE, 5X Denhardt’s solution (Invitrogen 750018), 0.1 mg/ml salmon sperm DNA (Agilent 201190), 1% SDS, 10% Dextran solution). Membrane was washed three times in 2X SSC/0.1% SDS (20X SSC: 3 M NaCl, 0.3 M Na Citrate, pH 7.0) and twice in 0.2X SSC/0.1% SDS, then exposed to a phosphor screen and developed using a phosphorimager (GE Healthcare Typhoon).

### Chromatin immunoprecipitation and library preparation

Chromatin immunoprecipitation (ChIP) was performed, with slight modification, as described by Thomas *et al.* (2015). Cells were first treated with 20nM 4-OHT for 24 hours, then crosslinked in 1% methanol-free FA (Thermo Fisher 28908) for 10 minutes and quenched using 0.125mM glycine. The cells were then rinsed twice in ice-cold 1X PBS, and lysed in formaldehyde lysis buffer (FALB: 50 mM HEPES pH 7.5, 140 mM NaCl, 1mM EDTA, and 1% Triton X-100) + 1% SDS + PIC (Roche 05056489001). Sonication was performed in a BioRuptor (Diagenode, Denville, New Jersey) for 25 minutes, 30s on/30s off, and debris removed by centrifugation. To enable ChIP efficiency to be determined by qPCR, a 1:50 (2%) sample of chromatin was removed (input) prior to antibody addition.

For ChIP, antibody (anti-HA, anti-IgG or anti-HCF1_N_) was added to chromatin prepared from 12×10^6^ Ramos cells, and samples rotated overnight at 4°C. Protein A Agarose (Roche 11134515001), blocked with 10μg BSA, was added to each sample and rotated at 4°C for 2-4 hours. Washes were performed with Low Salt Wash Buffer (20 mM Tris pH 8.0, 150 mM NaCl, 2 mM EDTA, 1% Triton X-100), High Salt Wash Buffer (20 mM Tris pH 8.0, 500 mM NaCl, 2 mM EDTA, 1% Triton X-100), Lithium Chloride Wash Buffer (10 mM Tris pH 8.0, 250 mM LiCl, 1 mM EDTA, 1% Triton X-100), and twice with TE (10 mM Tris pH 8.0, 1 mM EDTA). Input and ChIP sample crosslinking were reversed at 65°C overnight in 50 μl TE + 200 mM NaCl + 0.1% SDS + 20-40 μg PK (Macherey-Nagel 740506). For sequencing, three independent ChIPs (anti-IgG from CRE-ER^T2^ cells and anti-HCF1_N_ from MYC-WT cells) performed using the same antibody with the same chromatin were combined and DNA was extracted with phenol:chloroform:isoamyl alcohol, and ethanol precipitated with glycogen (Roche 10901393001). Libraries were prepared using NEBNext Ultra II DNA Library Prep Kit for Illumina (NEB E7645S) and NEBNext Multiplex Oligos for Illumina (NEB Set 1 E7335 and Set 2 E7500S). Additional AMPure clean-ups at the start and the end of library preparation were included. Sequencing was carried out by Vanderbilt Technologies for Advanced Genomics using Illumina NextSeq 500, 75bp paired-end reads. The total number of sequencing reads for each replicate is shown in Supplementary File 3.

For qPCR, samples (either input or ChIP) were brought up to a final volume of 200 μl using TE. Each reaction was performed in a final volume of 15 μl, containing 2X SYBR FAST qPCR Master Mix (Kapa, Wilmington, Massachusetts, KK4602), 300 nM of each primer, and 2 μl of diluted sample. Three technical replicates were performed for each sample, and the mean Ct of these was used for calculating percent input. The mean Ct value for input was adjusted using the equation Ct(input)-log_2_(50). Percent input was then calculated using the equation 100 *2^(adjustedCt-Ct(ChIP)). Three biological replicates of ChIP-qPCR were performed using primers to amplify across the genes EXOSC5, UTP20, EIF2S1, EIF4G3, and HBB (β-Globin).

### *In vitro* binding assays

pSUMO-MYC WT-FLAG containing 6XHis- and FLAG-tagged full-length MYC (Thomas et al., 2015) was used for Q5 site-directed mutagenesis to substitute in the 4A (4A_F and 4A_R) and VP16 HBM (VP16 HBM_F and VP16 HBM_R) mutations. Rosetta cells (Millipore 70954) were transformed with these plasmids, grown overnight, and induced the following day for three hours with 1 mM isopropyl β-d-1-thiogalactopyranoside (IPTG) at 30°C. Resulting cell pellets were resuspended in Buffer A (100 mM NaH2PO4, 10 mM Tris, 6 M GuHCl, 10 mM imidazole) + PIC (Roche 05056489001). Cells were sonicated 3X at 25% power for 10s (Cole-Parmer GE 130PB-1), and debris removed by centrifugation. 150 μl bed volume Ni-NTA agarose (QIAGEN, Venlo, Netherlands, 30210) was added to the samples and rotated for two hours at 4°C. Beads were sequentially washed 1X with Buffer A, 1X with Buffer A/TI (1:3 Buffer A:Buffer TI), 1X with Buffer TI (25 mM Tris-HCl pH 6.8, 20 mM imidazole), and 1X with SUMO wash buffer (3 mM imidazole, 10% glycerol, 1X PBS, 2 mM DTT) + PIC (Roche 05056489001). Recombinant MYC was eluted from the beads using SUMO elution buffer (250 mM imidazole, 10% glycerol, 1X PBS, 2 mM DTT) + PIC (Roche 05056489001).

HCF-1_VIC_ (residues 1-380) from pCGT-HCF1_VIC_ (Thomas et al., 2016) was cloned into pT7-IRES His-N (Takara 3290) using BamHI-HF (NEB R3136) and SalI-HF (NEB R3138), and Q5 site-directed mutagenesis was used to add an N-terminal T7 tag using the primers T7-HCF1_F and T7-HCF1_R. pT7-IRES His-T7-HCF-1_VIC_ was *in vitro* transcribed/translated using the TnT Quick Coupled Transcription/Translation System (Promega, Madison, Wisconsin, L1171). Two milligrams recombinant MYC and 12 μl T7-HCF-1_VIC_ were rotated overnight at 4°C in Kischkel buffer + PIC (Roche 05056489001). Anti-FLAG M2 Affinity Gel (Sigma-Aldrich), blocked with 10 μg BSA, was added to each sample and rotated for 2 hours at 4°C. Beads were washed 4X in Kischkel buffer + 2 μg/ml Aprotinin (VWR, Radnor, Pennsylvania, 97062-752) + 1 μg/ml Pepstatin (VWR 97063-246) + 1 μg/ml Leupeptin (VWR 89146-578), and incubated in 1X Laemmli buffer for 5 minutes at 95°C.

### Electrophoretic mobility shift assays

MYC:MAX dimers were purified and prepared as described by Farina *et al.* (2004). pRSET-6XHis-MYC WT, a gift from Ernest Martinez, was used as a template for Q5 site directed mutagenesis to substitute in the 4A (4A_F and 4A_R) and VP16 HBM (VP16 HBM_F and VP16 HBM_R) mutations. The resulting plasmids, or pET-His-MAX, also from Ernest Martinez, were transformed into Rosetta cells (Millipore 70954), grown overnight, and induced the following day for three hours with 1mM IPTG at 30°C. Resulting bacterial cell pellets were washed 1X with ice-cold wash buffer (10 mM Tris pH 7.9, 100 mM NaCl, 1 mM EDTA), then resuspended in lysis buffer (20 mM HEPES pH 7.9, 500 mM NaCl, 10% glycerol, 0.1% NP-40, 10 mM BME, 1 mM PMSF) and sonicated. Centrifugation was used to separate out the insoluble (pellet) and soluble (supernatant) fractions. For MAX, the recombinant protein is present in the supernatant, and was then purified using Ni-NTA agarose (see below). For MYC samples, the supernatant was discarded, and the insoluble pellet was resuspended in E-buffer (50 mM HEPES pH 7.9, 5% glycerol, 1% NP-40, 10% Na-DOC, 0.5 mM BME), lysed using a Dounce homogenizer and B-pestle, and centrifuged. The pelleted inclusion bodies were lysed overnight in S-buffer (10 mM HEPES pH 7.9, 6 M GuHCl, 5 mM BME) by shaking at 25°C, and debris spun out by centrifugation.

Both the MAX supernatant and MYC supernatant from lysed inclusion bodies were adjusted to 5mM imidazole. MYC and MAX were then bound to 75 μl bed volume Ni-NTA agarose (QIAGEN 30210). Successive washes were performed for 5 minutes each at 4°C: 3X with S-buffer + 5 mM imidazole, 3X with BC500 (20 mM Tris pH 7.9, 20% glycerol, 500 mM KCl, 0.05% NP-40, 10 mM BME, 0.2 mM PMSF) + 7 M urea + 5 mM imidazole, 1X with BC100 (20 mM Tris pH 7.9, 20% glycerol, 100 mM KCl, 0.05% NP-40, 10 mM BME, 0.2 mM PMSF) + 7 M urea + 15 mM imidazole, 1X with BC100 + 7 M urea + 30 mM imidazole. Elution was performed using BC100 + 7 M urea + 300 mM imidazole. Concentration of recombinant MYC and MAX was determined by running samples out on a 12% acrylamide gel alongside a BSA standard, and staining using Coomassie stain (50% methanol, 10% acetic acid, 0.1% w/v Coomassie Brilliant Blue).

For renaturation, 1.5 μg recombinant MAX was combined with 15 μg recombinant MYC (1:3 molar ratio), and brought up to a final volume of 150 μl using BC100 + 7 M urea. Dialysis was performed using a “Slide-A-Lyzer” (Thermo Fisher 66383), with each dialysis step done for 2 hours stirring in the following solutions: BC500 + 0.1% NP-40 + 4 M urea at room temperature, BC500 + 0.1% NP-40 + 2 M urea at room temperature, BC500 + 0.1% NP-40 + 1 M urea at room temperature, BC500 + 0.1% NP-40 + 0.5 M urea at room temperature, BC500 at 4°C, and twice BC100 at 4°C. The product was centrifuged to remove debris, and BSA was added to a final concentration of 500 ng/μl.

Double-stranded labeled E-box probe (biotin group at the 3’end) and unlabeled competitors were prepared with dsDNA buffer (30 mM Tris pH 7.9, 200 mM KCl), and incubated at 95°C for 5 minutes. The E-box sequence used was 5′-GCTCAGGGACCACGTGGTCGGGGATC-3′ and the mutant E-box sequence used was 5′-GCTCAGGGACCA<u>GC</u>TGGTCGGGGATC-3′ (IDT). Double-stranded probe, and the specific and non-specific competitors were prepared by combining 25 μM of each strand in dsDNA buffer (30 mM Tris pH 7.9, 200 mM KCl). For the probe, the forward strand carried a 3’ biotin group. 20 fmol of labeled probe was bound to 0.55 pmol MYC:MAX or 0.06 pmol MAX:MAX dimers in the presence of 20 ng poly(dI-dC) (Thermo Fisher 20148E) in binding reaction buffer (15 mM Tris pH 7.9, 15% glycerol, 100 mM KCl, 0.15 mM EDTA, 0.075% NP-40, 7.5 mM BME, 375 ng/μl BSA) for 30 minutes at room temperature. For reactions involving unlabeled specific or non-specific competitor, these were included in the binding reaction at a 100-fold excess over the biotinylated probe. EMSA gel loading solution (Thermo Fisher 20148K) was added to each sample and these were loaded onto a pre-run 6% polyacrylamide gel in 0.5X TBE (45 mM Tris, 45 mM boric acid, 1 mM EDTA). The gel was transferred to Hybond-N+ nylon membrane (GE Healthcare RPN303B) in 0.5X TBE for 30 minutes at 100 V. The remainder of the protocol was performed using LightShift Chemiluminescent EMSA Kit (Thermo Fisher 20148) according to manufacturer’s instructions.

### RNA preparation, RT-qPCR, and RNA-Seq

Cell pellets were resuspended in 1 ml TRIzol (Invitrogen 15596026), and RNA was prepared according to the manufacturer’s instructions. For switchable MYC allele Ramos cells, cells were treated with 20 nM 4-OHT for 24 hours, and harvested. Prepared RNA was submitted to GENEWIZ (South Plainfield, NJ) for DNAse treatment, rRNA depletion, library preparation, and 150bp paired-end sequencing on Illumina HiSeq. For untagged or FKBP^FV^-HCF-1_N_ Ramos cells, cells were treated with DMSO or 500 nM dTAG-47 for 3 hours, and prepared RNA was DNAse treated prior to submission to Vanderbilt Technologies for Advanced Genomics for rRNA depletion, library preparation, and 150bp paired-end sequencing on Illumina NovaSeq 6000. The total number of sequencing reads for each replicate is shown in Supplementary File 3.

### Next generation sequencing analyses

After adapter trimming by Cutadapt (Martin, 2011), RNA-Seq reads were aligned to the hg19 genome using STAR (Dobin et al., 2013) and quantified by featureCounts (Liao et al., 2014). Differential analysis were performed by DESeq2 (Love et al., 2014), which estimated the log2 fold changes, Wald test P-values, and adjusted P-value (false discovery rate, FDR) by the Benjamini-Hochberg procedure. The significantly changed genes were chosen with the criteria FDR < 0.05. ChIP-Seq reads were aligned to the hg19 genome using Bowtie2 (Langmead et al., 2009) after adapter trimming. Peaks were called by MACS2 (Feng et al., 2012) with a q-value of 0.01. ChIP read counts were calculated using DiffBind (Stark and Brown, 2011). Peaks were annotated using Homer command annotatePeaks, and enriched motifs were identified by Homer command findMotifsGenome (http://homer.ucsd.edu/homer/).

### Cell growth and glutamine deprivation assays

To measure the impact of HCF-1_N_ degradation on cell growth, FKBP^FV^-HCF-1_N_ Ramos cells were plated at a density of 20,000 cells/ml with either DMSO or 500 nM dTAG-47. Cells were then counted every 24 hours for the following four days, without replacement of the compound or changing of the media.

To measure the impact of altering the MYC−HCF-1 interaction on cell growth, switchable MYC allele Ramos cell lines were treated with 20 nM 4-OHT for 2 hours to create a 50/50 mix of GFP-negative (WT) to GFP-positive (mutant) cells. Cells were sampled 24 hours later, and every three days following. Sampled cells were fixed in 1% FA in PBS for 10 minutes at room temperature, and the proportion of GFP-positive cells was determined using a BD LSR II flow cytometer at the Vanderbilt Flow Cytometry Shared Resource. To account for variation in the proportion of GFP-positive cells between replicates, each replicate was normalized to the proportion of GFP-positive cells at 24 hours post-treatment with 4-OHT.

To measure the impact of altering the MYC−HCF-1 interaction on glutamine dependence, switchable MYC allele Ramos cell lines were treated with 20 nM 4-OHT for 2 hours, and allowed to recover for three days. Cells were then split into RPMI 1640 without L-Glutamine (Corning 15-040-CV), supplemented with 10% dialyzed FBS (Gemini Bio, West Sacramento, California, 100-108), and 1% P/S (Gibco 15140122), and grown for 16 hours with or without supplemental 2mM glutamine (Gibco 25030081). Glutamine was added back to the cells that were deprived, grown for three days, and fixed in 1% FA in PBS for 10 minutes at room temperature. The proportion of GFP-positive cells was determined using a BD LSR II flow cytometer, and normalized to the proportion of GFP-positive cells prior to being grown with or without supplemental glutamine. For the 4A and VP16 HBM cells, the proportion of GFP-positive cells was normalized to that in WT for glutamine supplementation (Gln+) or deprivation (Gln-).

### Metabolomics

#### Sample Preparation

Global, untargeted metabolomics was performed on switchable MYC allele Ramos cell lines treated with 20 nM 4-OHT for 24 hours. Individual cell pellet samples were lysed using 200 μl ice cold lysis buffer (1:1:2, Acetonitrile : Methanol : Ammonium Bicarbonate 0.1 M, pH 8.0, LC-MS grade) and sonicated using a probe tip sonicator, 10 pulses at 30% power, cooling down on ice between samples. A bicinchoninic acid protein assay was used to determine the protein concentration for individual samples, and adjusted to 200 μg total protein in 200 μl of lysis buffer. Isotopically labeled standard molecules, Phenylalanine-D8 and Biotin-D2, were added to each sample to assess sample extraction quality. Samples were subjected to protein precipitation by addition of 800 μl of ice cold methanol (4X by volume), and incubated at −80°C overnight. Samples were centrifuged at 10,000 rpm for 10 minutes to eliminate precipitated proteins and supernatant(s) were transferred to a clean microcentrifugre tube and dried down *in vacuo*. Samples were stored at −80°C prior to LC-MS analysis.

#### Global untargeted LC-MS/MS analysis

For mass spectrometry analysis, individual samples were reconstituted in 50 μl of appropriate reconstitution buffer (HILIC: acetonitrile/ H_2_O, 90:10, v/v, RPLC: acetonitrile/H_2_O with 0.1% formic acid, 3:97, v/v). Samples were vortexed well to solubilize the metabolites and cleared by centrifugation using a benchtop mini centrifuge to remove insoluble material. Quality control samples were prepared by pooling equal volumes from each sample. During final reconstitution, isotopically labeled standard molecules, Tryptophan-D3, Carnitine-D9, Valine-D8, and Inosine-4N15, were spiked into each sample to assess LC-MS instrument performance and ionization efficiency.

High resolution (HR) MS and data-dependent acquisition (MS/MS) analyses were performed on a high resolution Q-Exactive HF hybrid quadrupole-Orbitrap mass spectrometer (Thermo Fisher Scientific, Bremen, Germany) equipped with a Vanquish UHPLC binary system and autosampler (Thermo Fisher Scientific, Germany) at the Vanderbilt Center for Innovative Technology. For HILIC analysis metabolite extracts (10 μl injection volume) were separated on a SeQuant ZIC-HILIC 3.5-μm, 2.1 mm × 100 mm column (Millipore Corporation, Darmstadt, Germany) held at 40°C. Liquid chromatography was performed at a 200 μl min^−1^ using solvent A (5 mM Ammonium formate in 90% H_2_O, 10% acetonitrile) and solvent B (5 mM Ammonium formate in 90% acetonitrile, 10% H_2_O) with the following gradient: 95% B for 2 min, 95-40% B over 16 min, 40% B held 2 min, and 40-95% B over 15 min, 95% B held 10 min (gradient length 45 min). For the RPLC analysis metabolite extracts (10 μl injection volume) were separated on a Hypersil Gold, 1.9 μm, 2.1 mm x 100 mm column (Thermo Fisher) held at 40°C. Liquid chromatography was performed at a 250 μl/min using solvent A (0.1% formic acid in H_2_O) and solvent B (0.1% formic acid in acetonitrile) with the following gradient: 5% B for 1 minute, 5-50% B over 9 minutes, 50-70% B over 5 minutes, 70-95% B over 5 minutes, 95% B held 2 minutes, and 95-5% B over 3 minutes, 5% B held 5 minutes (gradient length: 30 minutes).

Full MS analyses were acquired over a mass range of m/z 70-1050 using electrospray ionization positive mode. Full mass scan was used at a resolution of 120,000 with a scan rate of 3.5 Hz. The automatic gain control (AGC) target was set at 1 × 10^6^ ions, and maximum ion injection time was at 100 ms. Source ionization parameters were optimized with the spray voltage at 3.0 kV, and other parameters were as follows: transfer temperature at 280 °C; S-Lens level at 40; heater temperature at 325°C; Sheath gas at 40, Aux gas at 10, and sweep gas flow at 1.

Tandem mass spectra were acquired using a data dependent scanning mode in which one full MS scan (m/z 70-1050) was followed by 2, 4 or 6 MS/MS scans. MS/MS scans were acquired in profile mode using an isolation width of 1.3 m/z, stepped collision energy (NCE 20, 40), and a dynamic exclusion of 6 s. MS/MS spectra were collected at a resolution of 15000, with an AGC target set at 2 × 10^5^ ions, and maximum ion injection time of 100 ms. The retention times and peak areas of the isotopically labeled standards were used to assess data quality.

#### Metabolite data processing and analysis

LC-HR MS/MS raw data were imported, processed, normalized and reviewed using Progenesis QI v.2.1 (Non-linear Dynamics, Newcastle, UK). All MS and MS/MS sample runs were aligned against a quality control (pooled) reference run, and peak picking was performed on individual aligned runs to create an aggregate data set. Unique ions (retention time and m/z pairs) were grouped (a sum of the abundancies of unique ions) using both adduct and isotope deconvolutions to generate unique “features” (retention time and m/z pairs) representative of unannotated metabolites. Data were normalized to all features using Progenesis QI. Compounds with <25% coefficient of variance (%CV) were retained for further analysis. Variance stabilized measurements achieved through log normalization were used with Progenesis QI to calculate P-values by one-way analysis of variance (ANOVA) test and adjusted P-values (false discovery rate, FDR). Significantly changed metabolites were chosen with the criteria FDR < 0.05 & |FC| > 1.5.

Tentative and putative identifications were determined within Progenesis QI using accurate mass measurements (<5 ppm error), isotope distribution similarity, and fragmentation spectrum matching based on database searches against Human Metabolome Database (HMDB) (Wishart et al., 2013), METLIN (Smith et al., 2005), the National Institute of Standards and Technology (NIST) database (Salvat et al., 2016), and an in-house library. In these experiments, the level system for metabolite identification confidence was utilized (Schrimpe-Rutledge et al., 2016). Briefly, many annotations were considered to be tentative (level 3, L3) and/or putative (level 2, L2); in numerous circumstances a top candidate cannot be prioritized, thus annotations may represent families of molecules that cannot be distinguished.

### *In vivo* studies

#### Tumor formation and maintenance assays

Six-week-old athymic nude mice (female *Foxn1^nu/nu^*; Envigo, Indianapolis, IN) were injected subcutaneously into one flank with 10^7^ switched or unswitched WT, Δ264, 4A-1 or 4A-2 cells at the Thomas Jefferson Research Animals Shared Resource Core. 4A-1 and 4A-2 carry the same 4A mutation, but were independent clones obtained from the same population. To facilitate lymphoma cell seeding, mice received whole-body irradiation (6 Gy) 24 hours prior to cell injection. For tumor formation studies, lymphoma cells were treated *in vitro* with 4-OHT for 24 hours to induce the switchable MYC cassette and expanded for two days prior to being injected into mice. For tumor maintenance studies, mice were injected with unswitched cells and allowed to form palpable tumors. Once tumors reached approximately 200 mm^3^, mice received intraperitoneal injections of tamoxifen (2 mg in corn oil, Sigma-Aldrich T5648) once daily for 3 consecutive days to induce the switchable MYC cassette *in vivo*. Digital calipers were used to measure tumors and volumes calculated using the ellipsoid formula. Mice were sacrificed at humane endpoints based on tumor volume. Kaplan-Meier survival analyses were compared by log-rank tests to determine statistical significance. For lymphoma cell apoptosis evaluation, a cohort of mice with size-matched tumors prior to tamoxifen injection were sacrificed 48 and 96 hours following the first administration of tamoxifen. Flow cytometry (BD LSRII) at Thomas Jefferson University Flow Cytometry Shared Resource was used to measure fragmented (subG1) apoptotic DNA with propidium iodide (Sigma-Aldrich P4170), Annexin V/7AAD (BD Pharmingen 559763), and Caspase 3 activity (BD Pharmingen) in the lymphoma cells isolated from the tumors, as we previously reported (Adams et al., 2017) and as per manufacturer’s protocols. Two-tailed t-tests were used to determine significance when comparing two groups. All mouse experiments were approved by the Institutional Animal Care and Use Committee at Thomas Jefferson University and complied with state and federal guidelines.

#### Tumor gDNA Analysis

To determine the proportion of cells that remain switched in the resulting tumors, gDNA was prepared from tumors using the PureLink Genomic DNA Mini Kit (Invitrogen K182002) according to manufacturer’s instructions. As described in Thomas *et al.* (2019), gDNA from 0% switched (switchable MYC WT cells grown in puromycin) and 100% switched (permanently switched clonal cells) was also prepared in this manner for normalization. The resulting gDNA was diluted down to 50 ng/μl, so that 2 μl (100 ng) was loaded per well for qPCR. qPCR was performed using 2X SYBR FAST qPCR Master Mix (Kapa, Wilmington, Massachusetts, KK4602) in a Bio-Rad CFX96 Real-Time System. For each primer set (MYCP-4 and MYCP-5; SNHG15_F and SNHG15_R) and each independent tumor replicate, an average of three qPCR wells (technical replicates) was used. MYCP-4 and MYCP-5 primer set only amplifies gDNA from unswitched cells, whereas SNHG15_F and SNHG15_R amplifies gDNA from both unswitched and switched cells. For each gDNA sample, ΔCt was calculated as the difference between MYCP and SNHG15. gDNA from the 0% and 100% switched cells was used to calculate ΔΔCt, which was then used to normalize the ΔCt for the tumor gDNA to estimate the proportion of switched cells.

#### Tumor RNA-Seq

Tumors from mice sacrificed 48 hours after the first tamoxifen administration were submitted to GENEWIZ for RNA extraction, DNAse treatment, rRNA depletion, library preparation, and 150bp paired-end sequencing on Illumina HiSeq. Four tumors for each WT, 4A-1, 4A-2, and Δ264 were submitted. The total number of sequencing reads for each replicate is shown in Supplementary File 3.

### Pathway and gene ontology analysis, and figure generation

Classification of annotated metabolites was extracted from HMDB (Wishart et al., 2018) and LIPID MAPS (Fahy et al., 2009). Metabolite pathway analyses were performed using MetaboAnalyst 4.0 (Chong et al., 2019), and gene ontology (GO) analyses using Metascape (Zhou et al., 2019). Chord diagrams were created using Circos (Krzywinski et al., 2009), pathways using Cytoscape (Shannon et al., 2003), and bubble plots using ggplot2 (Wickham, 2016).

### Statistical analysis and replicates for non-high throughput data

Unless otherwise stated, all experiments were conducted with at least three independent, biological replicates, and statistical tests were carried out using PRISM 8 (GraphPad).

## ACKNOWLEDGEMENTS

We thank Winship Herr for comments on the manuscript. For reagents we thank James Bradner, Feng Zhang, and Ernest Martinez. We thank the Vanderbilt Institute of Chemical Biology Synthesis Core for synthesis of dTAG-47. The VU Flow Cytometry Shared Resource is supported by the Vanderbilt Ingram Cancer Center (P30CA68485) and the Vanderbilt Digestive Disease Research Center (DK058404). VANTAGE is supported by the Vanderbilt Ingram Cancer Center, the Vanderbilt Digestive Disease Research Center, and the Vanderbilt Vision Center (P30EY08126). The Thomas Jefferson Flow Cytometry and Research Animals shared resource cores are supported by the Sidney Kimmel Cancer Center (NCI/NIH P30CA056036). This work was supported in part using the resources of the Center for Innovative Technology at Vanderbilt University. This work was supported by the Vanderbilt International Scholars Program (TMP), Robert J. Kleberg, Jr., and Helen C. Kleberg Foundation (WPT), Edward P. Evans Foundation (WPT), the NCI/NIH (CA200709; WPT), the NCI/NIH (CA211305; LRT), the NCI/NIH (CA148950 CME), the Integrated Biological Systems Training in Oncology Training Program (T32 CA119925; AMW), the Rally Foundation for Childhood Cancer Research Fellowship (AMW), Open Hands Overflowing Hearts co-funded research fellowship (AMW), and the American Association for Cancer Research Basic Cancer Research Fellowship (AMW), the Herbert A. Rosenthal, MD’56 endowed professorship in Cancer Research (CME), the Steinfort family fund (CME), and the Sidney Kimmel Cancer Center/Thomas Jefferson University (CME, CMA).

## SUPPLEMENTAL INFORMATION INDEX

Figure 1 - figure supplement 1: Validation of switchable Ramos cell lines to study the MYC–HCF-1 interaction.

Figure 2 - figure supplement 1: Impact of the 4A and VP16 HBM mutants on Ramos cell metabolism.

Figure 2 - Source Data 1: Ramos 4A and VP16 HBM RPLC significant changes

Figure 2 - Source Data 2: Ramos 4A and VP16 HBM HILIC significant changes

Figure 3 - figure supplement 1: Gene expression changes induced by the 4A and VP16 HBM mutants.

Figure 3 - figure supplement 2: Amino acids and their cognate tRNA-ligases are influenced by the MYC– HCF-1 interaction.

Figure 3 - Source Data 1: Ramos 4A 24 hour RNA-Seq significant changes

Figure 3 - Source Data 2: Ramos VP16 HBM 24 hour RNA-Seq significant changes

Figure 3 - Source Data 3: Ramos 4A and VP16 HBM 24 hour RNA-Seq shared significant changes

Figure 4 - figure supplement 1: Inducible degradation of HCF-1_N_.

Figure 4 - Source Data 1: Ramos HCF-1_N_ degradation RNA-Seq significant changes

Figure 4 - Source Data 2: Ramos untagged RNA-Seq significant changes

Figure 5 - figure supplement 1: Comparison of MYC and HCF-1_N_ localization on Ramos cell chromatin.

Figure 5 - figure supplement 2: HBM mutations do not impact DNA-binding by MYC:MAX complexes.

Figure 5 - Source Data 1: Ramos HCF-1_N_ annotated ChIP-Seq peaks

Figure 5 - Source Data 2: Annotated intersect of ChIP-Seq peaks for HCF-1_N_ and MYC in Ramos cells

Figure 6 - figure supplement 1: Disruption of the MYC−HCF-1 interaction *in vivo*.

Figure 6 - Source Data 1: Tumor RNA-Seq significant changes

Supplementary File 1: Primer sequences

Supplementary File 2: Example flow cytometry gating

Supplementary File 3: Next-generation sequencing read counts

---

**Figure 1 - figure supplement 1:**
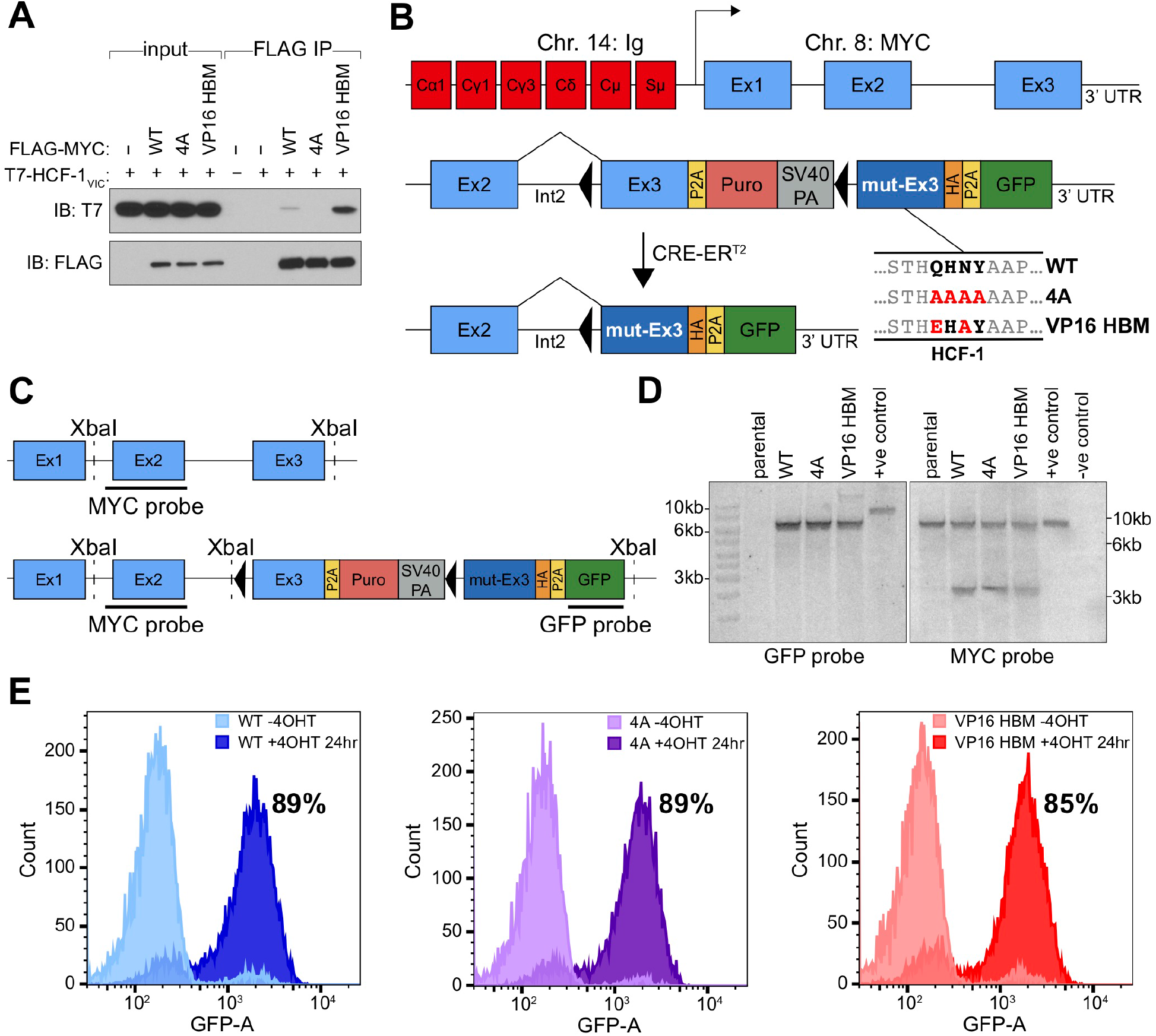
Validation of switchable Ramos cell lines to study the MYC–HCF-1 interaction. (**A**) *In vitro* transcribed/translated T7-tagged HCF-1_VIC_ was incubated with recombinant FLAG-tagged c-MYC, either WT or mutant (4A or VP16 HBM), and immunoprecipitation (IP) performed using anti-FLAG M2 agarose. Western blot of the input lysate, and the IP eluate, was performed using antibodies against the T7 and FLAG tags. (**B**) The translocated *MYC* locus from Ramos cells is depicted at top, with chromosome 14 (red) and 8 (blue) elements indicated. Beneath is a representation of the locus modification, in either the unswitched (middle) or switched (bottom) states. This switchable allele contains a wild-type (WT) exon 3, a P2A-linked puromycin cassette, and a SV40 polyadenylation (SV40 PA) signal, all of which are flanked by LoxP sites (black triangles). Downstream of the LoxP-flanked region is an HA-tagged mutant exon 3 (mut-Ex3) and a P2A-linked GFP cassette. Activation of CRE-ER^T2^ results in excision of WT exon 3 and its replacement with mutant exon 3 which carries sequences encoding either WT or mutant (4A or VP16 HBM) MYC protein. (**C**) Comparison of the structure of the parental (non-modified) *MYC* allele (top) compared to the switchable *MYC* allele (bottom). XbaI sites used for digestion of genomic DNA in Southern blot are highlighted, as are the complementary sites for the MYC and GFP probes. The expected products of XbaI digestion for the parental line are 6,784 bp for the MYC probe (which detects both the translocated and non-translocated alleles) and nothing for the GFP probe; for correctly-engineered lines the expected sizes are 6,784 bp and 2,942 bp for the MYC probe, and 6,624 bp for the GFP probe. (**D**) Southern blot using GFP and MYC probes on XbaI-digested gDNA from unswitched parental or switchable cells (WT, 4A, and VP16 HBM), with digested positive and negative control plasmids. (**E**) Switchable cells were treated with or without 20 nM 4-OHT (24 hours), fixed using 1% formaldehyde, and subject to flow cytometry. The GFP profiles of the −4-OHT and +4-OHT cells are shown overlaid onto the same axes, with the approximate percentage of GFP-positive cells for 24 hours +4-OHT shown.

**Figure 2 - figure supplement 1:**
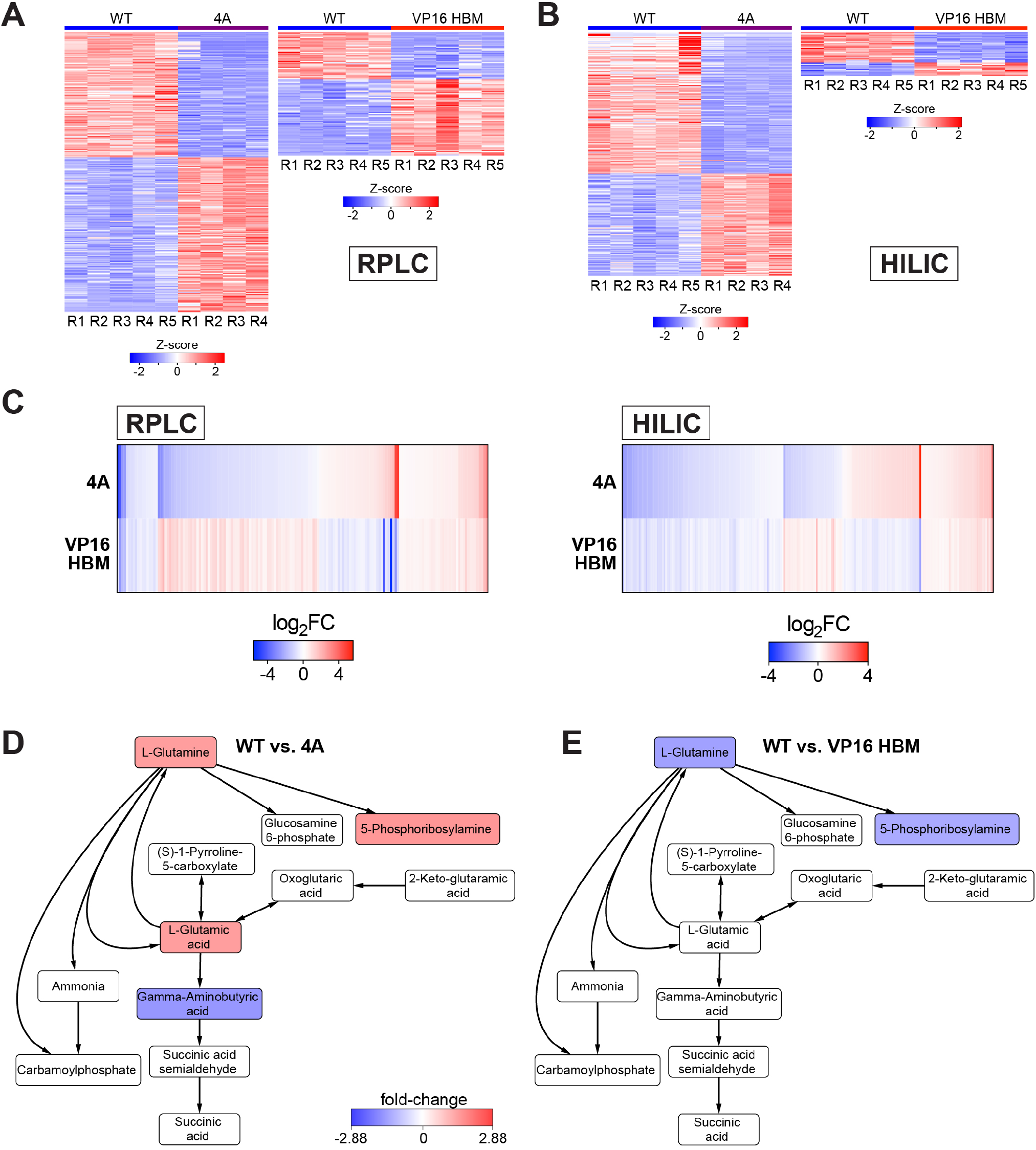
Impact of the 4A and VP16 HBM mutants on Ramos cell metabolism. All data are derived from switchable Ramos cells treated with 20 nM 4-OHT for 24 hours. (**A**) Heatmap representing metabolites (z-transformed) detected with RPLC that were significantly (FDR < 0.05 & |FC| > 1.5) impacted with the 4A (left) or VP16 HBM (right) MYC mutants compared to WT. (**B**) As in (A) but for metabolites detected by HILIC. (**C**) Heatmap showing metabolites detected with RPLC (left) or HILIC (right) that are significantly (FDR < 0.05) changed for both the 4A and VP16 HBM mutants, compared to WT MYC. Metabolites are clustered according to the relationship between the two mutants, and ranked by the log_2_FC of 4A versus WT. (**D**) Metabolites in the “alanine, aspartate, and glutamate metabolism” pathway that are significantly (FDR < 0.05) impacted in the 4A mutant. Node color represents the fold-change over WT. The remainder of pathway is shown in **Figure 2J**. (**E**) Metabolites in the “alanine, aspartate, and glutamate metabolism” pathway that are significantly (FDR < 0.05) impacted in the VP16 HBM mutant. Node color represents the fold-change over WT. The remainder of pathway is shown in **Figure 2J**.

**Figure 3 - figure supplement 1:**
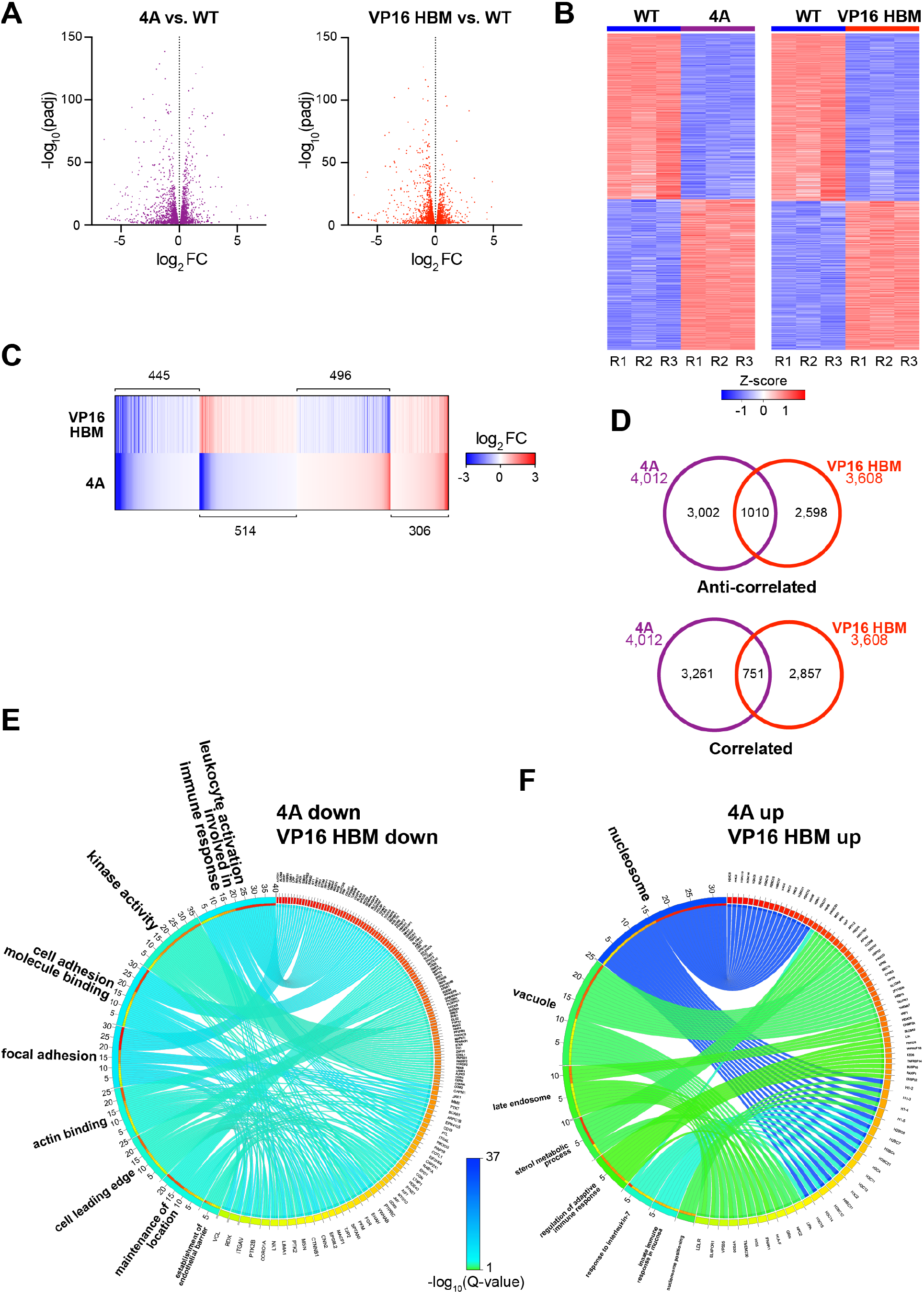
Gene expression changes induced by the 4A and VP16 HBM mutants. (**A**) Volcano plots showing the significant (FDR < 0.05) gene expression changes for the 4A (left) and VP16 HBM (right) mutants. For clarity, some data points were excluded; these data points are highlighted in Figure 3 - Source Data 1 and 2. (**B**) Heatmap showing z-transformed RNA-Seq data for switchable MYC allele cell lines at 24 hours, ranked by FC. Genes that were significantly (FDR < 0.05) impacted compared to WT are shown. Three biological replicates for each WT, 4A, and VP16 HBM were used to calculate FDR and fold-changes. (**C**) Heatmap showing the log_2_FC of significantly (FDR < 0.05) changed genes that are shared between the 4A and VP16 HBM mutants. Genes are clustered according to the relationship in expression changes between the 4A and VP16 HBM mutants, and ranked by the log_2_FC for the 4A mutant. Scale of heatmap is limited to [−3,3]. (**D**) Significant (FDR < 0.05) gene expression changes that are anti-correlative (top) and correlative (bottom) in direction between the 4A and VP16 HBM mutants. (**E**) and (**F**) Categories from the top eight families in GO enrichment analysis of the correlative gene clusters shown in (C), for genes that were decreased (E) or increased (F) with both mutants. The Q-value of categories is represented by the ribbon color, which is scaled across these figure and **Figures 3D and 3E**. Categories are ranked by the number of matched genes, and genes are ranked by the number of categories into which they fall.

**Figure 3 - figure supplement 2:**
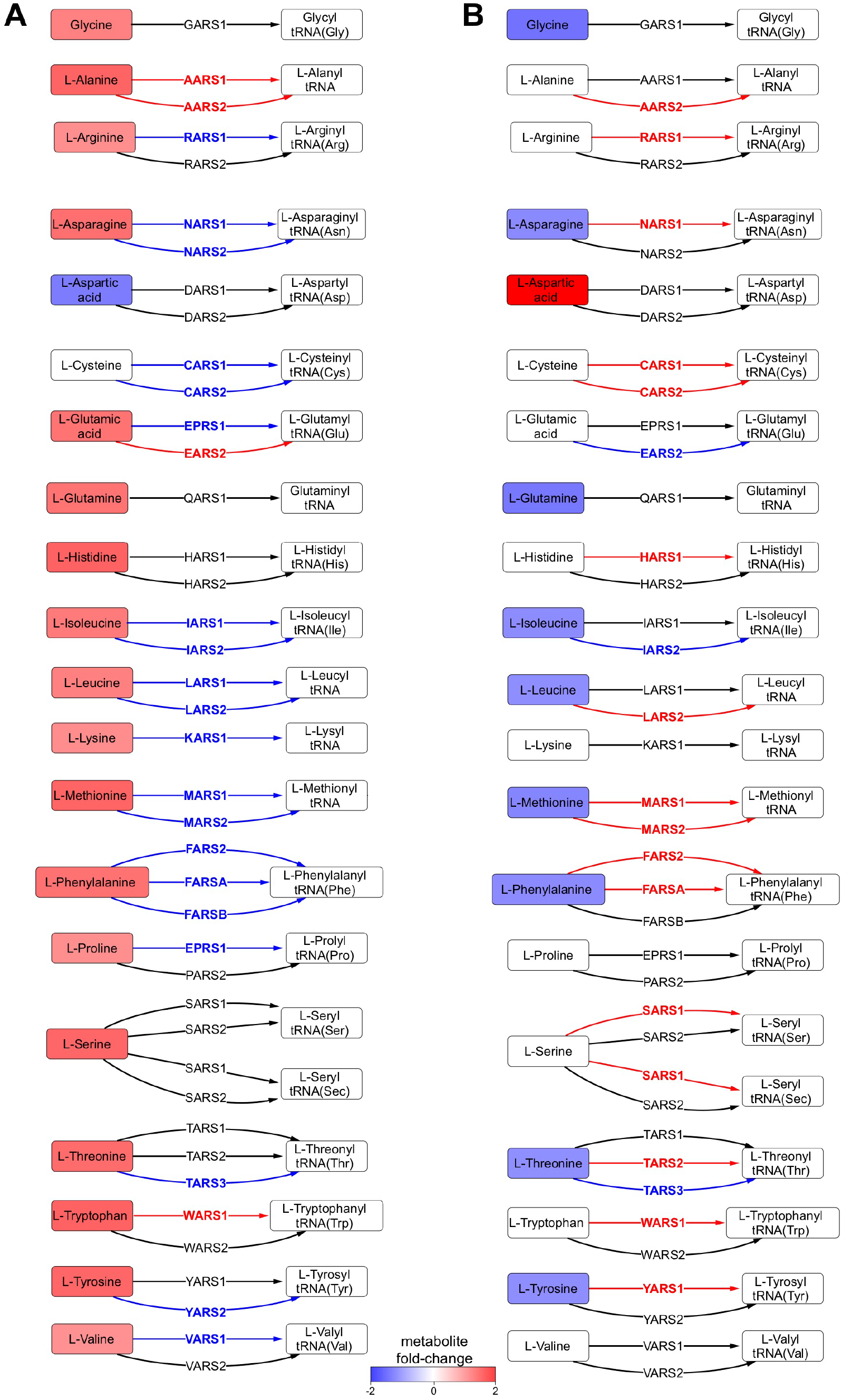
Amino acids and their cognate tRNA-ligases are influenced by the MYC–HCF-1 interaction. Impact of the 4A (**A**) and VP16 HBM (**B**) mutants on amino acid levels (HILIC) and on the expression of aminoacyl tRNA transferases (RNA-Seq). Genes and the reactions they catalyze are highlighted according the direction of RNA changes against WT, with blue reflecting decreases and red reflecting increases.

**Figure 4 - figure supplement 1:**
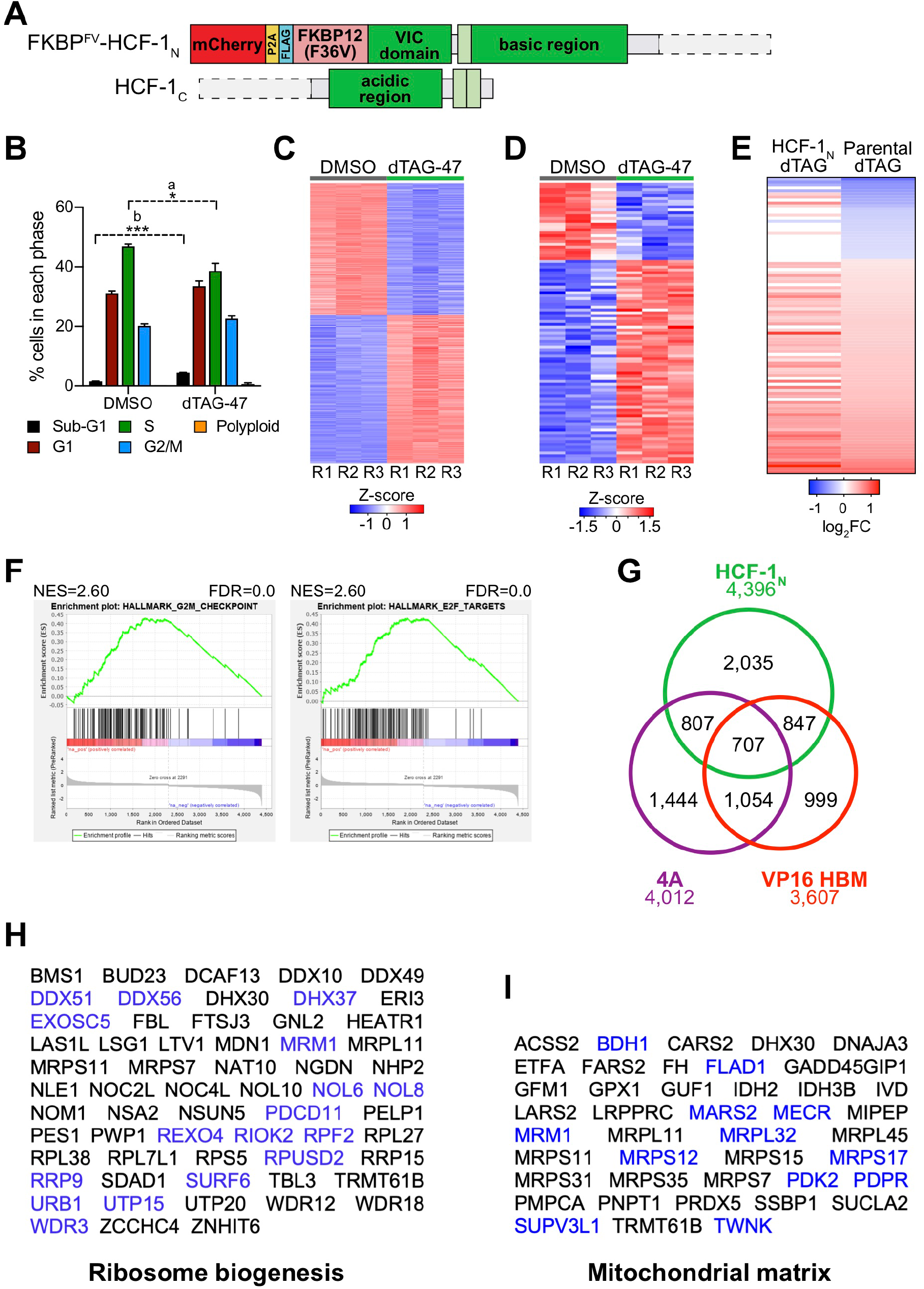
Inducible degradation of HCF-1_N_. (**A**) Schematic of the HCF-1 fusion protein in Ramos FKBP^FV^-HCF-1_N_ Ramos cells, which were generated using CRISPR/Cas9 homology-directed repair. Fused to the N-terminus of the VP16 induced complex (VIC) domain is mCherry linked by P2A to FLAG-tagged FKBP12^FV^. (**B**) Cell cycle distribution was determined using propidium iodide (PI) staining of FKBP^FV^-HCF-1_N_ Ramos cells treated with DMSO or 500 nM dTAG-47 for 24 hours. Shown are the mean and standard error for three biological replicates. Student’s t-test was used to calculate P-values; a = 0.037, b < 0.0001. (**C**) Heatmap of significantly (FDR < 0.05) changed genes in FKBP^FV^-HCF-1_N_ Ramos cells treated with DMSO or dTAG-47 three hours. Shown are the z-transformed expression data for three biological replicates of RNA-Seq, ranked by FC. (**D**) Heatmap of significantly (FDR < 0.05) changed genes in parental Ramos cells treated with DMSO or 500 nM dTAG-47 for three hours. Shown are the z-transformed expression data for three biological replicates of RNA-Seq, ranked by FC. (**E**) Heatmap showing the log_2_FC of all genes that are significantly (FDR < 0.05) changed when untagged Ramos cells were treated with DMSO or dTAG-47 for three hours, and the corresponding log_2_FC data in FKBP^FV^-HCF-1_N_ Ramos cells. (**F**) Gene set enrichment analysis (GSEA) of significantly (FDR < 0.05) changed genes in FKBP^FV^-HCF-1_N_ Ramos cells. Shown are the normalized enrichment scores (NES), and FDR for each gene set. (**G**) Venn diagram showing overlap of gene transcripts that are significantly (FDR < 0.05) changed in RNA-Seq with the 4A and VP16 HBM mutation, and with degradation of HCF-1_N_ in Ramos cells. (**H**) and (**I**) Genes that are decreased with the 4A MYC mutant and increased with the VP16 HBM MYC mutant and fall into the “Ribosome biogenesis” (H) or “Mitochondrial matrix” (I) gene ontology categories. Highlighted in blue are the genes that are also significantly decreased with degradation of HCF-1_N_.

**Figure 5 - figure supplement 1:**
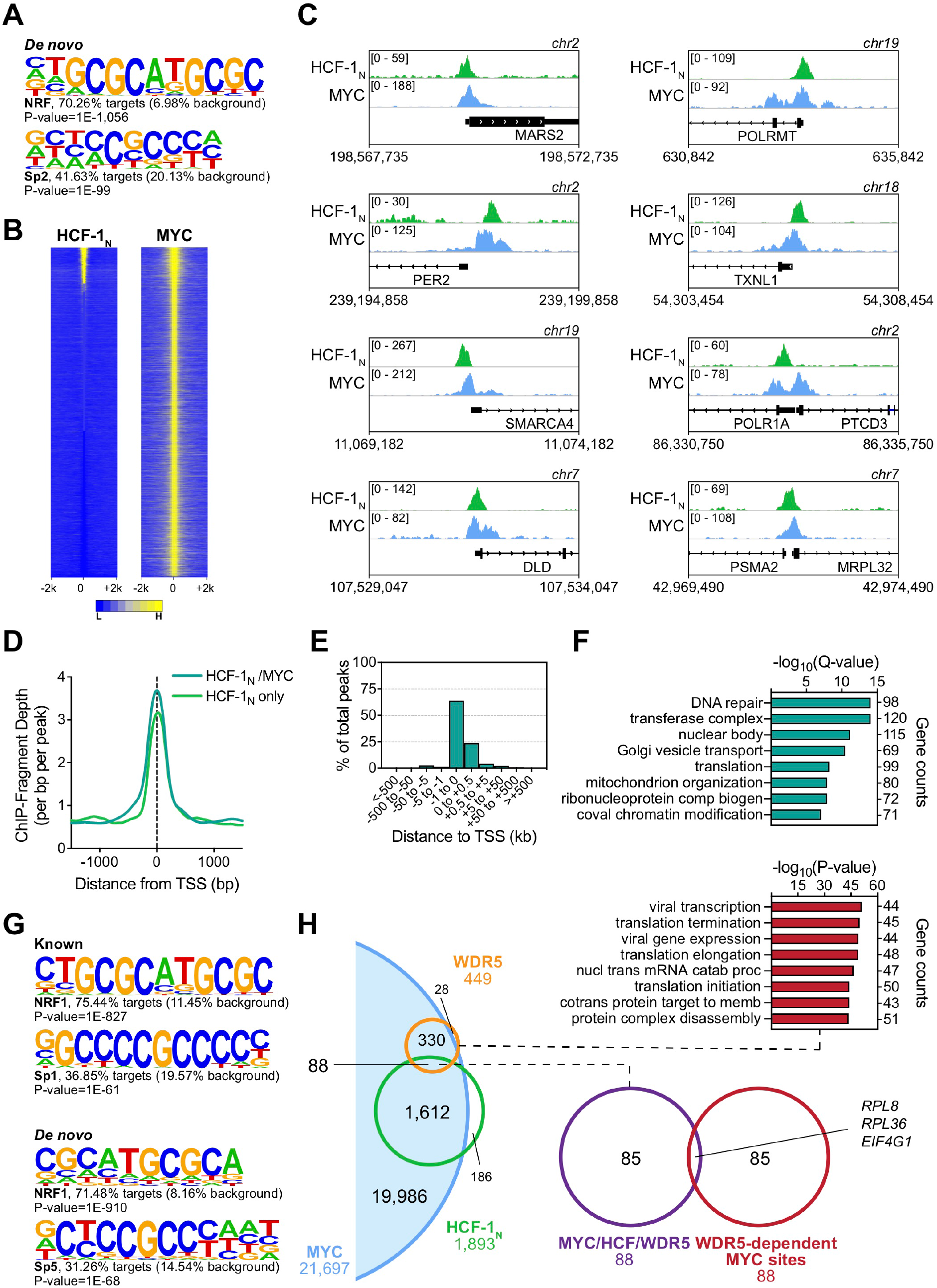
Comparison of MYC and HCF-1_N_ localization on Ramos cell chromatin. (**A**) *De novo* motif analysis of HCF-1_N_ peaks in Ramos cells. Two of the most highly enriched motifs are shown, as well the percentage of target and background sequences with the motif, and the P-value. (**B**) Heatmap of all MYC peaks in Ramos cells (from GSE126207) and the corresponding region in Ramos HCF-1_N_ ChIP-Seq, representing the combined average of normalized peak intensity in ± 2 kb regions surrounding the peak centers with 100-bp bin sizes. Ranking is by peak intensity in HCF-1_N_. (**C**) Example IGV screenshots of regions that have overlapping MYC and HCF-1_N_ in Ramos cells. (**D**) Normalized HCF-1_N_ ChIP-Seq fragment counts where peaks overlap with MYC (HCF-1_N_/MYC) compared to where they do not overlap (HCF-1_N_ only). Data are smoothed with a cubic spline transformation. (**E**) Distribution of regions that overlap between MYC and HCF-1_N_ peaks in Ramos cells in relation to the nearest TSS. (**F**) GO enrichment analysis of genes nearest to regions of overlap between MYC and HCF-1_N_ peaks. (**G**) Known and *de novo* motif analysis of regions that overlap between MYC and HCF-1_N_ peaks in Ramos cells. For each motif represented, percentage of target and background sequences with the motif, and P-value are shown. (**H**) Venn diagram showing relationship between HCF-1_N_, MYC, and WDR5 peaks in Ramos cells, and the overlap between co-bound genes and genes for which WDR5 is responsible for MYC recruitment. GO enrichment analysis of genes co-bound by MYC and WDR5—taken from Thomas *et al.* (2019)—is also shown

**Figure 5 - figure supplement 2:**
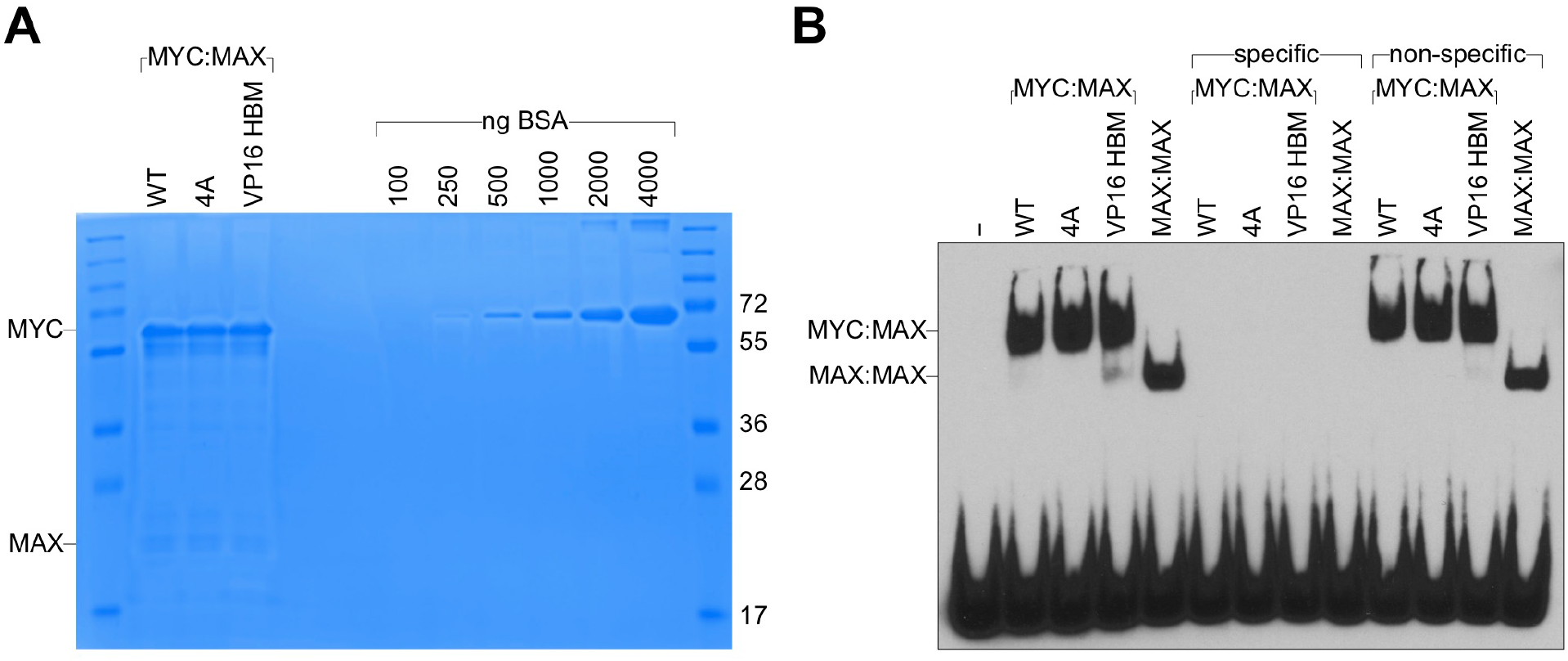
HBM mutations do not impact DNA-binding by MYC:MAX complexes. (**A**) Recombinant MYC (WT, 4A, or VP16 HBM):MAX dimers used in these assays. Dimers were separated by SDS-PAGE alongside a BSA standard, and stained using Coomassie Brilliant Blue. (**B**) Electrophoretic mobility shift assay (EMSA) using recombinant MYC:MAX and MAX:MAX dimers incubated with a biotinylated E-box probe (5′-GCTCAGGGACCACGTGGTCGGGGATC-3′). Binding reactions were conducted with or without 100-fold excess of unlabeled specific (as above) or non-specific (5′-GCTCAGGGACCA<u>GC</u>TGGTCGGGGATC-3′) competitors.

**Figure 6 - figure supplement 1:**
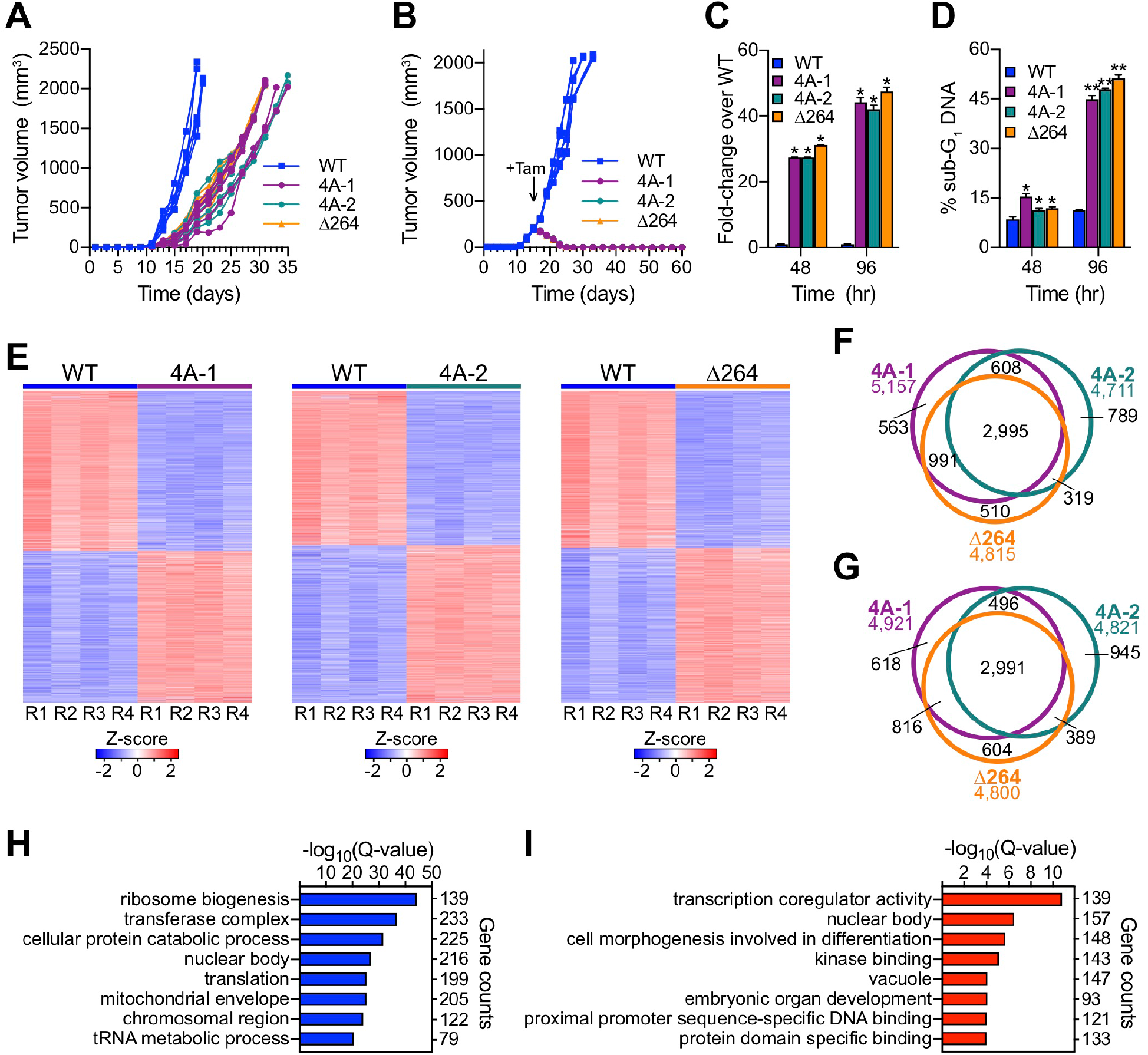
Disruption of the MYC−HCF-1 interaction *in vivo*. (**A**) Tumor volumes for individual mice in the tumor engraftment assay. (**B**) Tumor volumes for individual mice in the tumor maintenance assay. The day at which tamoxifen (Tam) administration was initiated is indicated with an arrow. (**C**) Caspase activity in cells isolated from tumors (n=4 each) at 48 and 96 hours following the first tamoxifen administration was measured. Shown are the mean and standard error for the fold-change (FC) over WT for each. Student’s t-test between WT and each of the mutants was used to calculate P-value; *P < 0.000001. (**D**) Propidium staining and flow cytometry were performed on cells isolated from four tumors each at 48 and 96 hours following the first tamoxifen administration to determine the extent of cell death. Shown are the mean and standard error. Student’s t-test between WT and each of the mutants was used to calculate P-value; *P < 0.000628, **P < 0.000001. (**E**) z-transformed RNA-Seq data from tumors (n=4) extracted after switching, ranked by FC. Genes that were significantly (FDR < 0.05) impacted compared to WT are shown. (**F**) Venn diagram owing the overlap of genes significantly (FDR < 0.05) decreased in the 4A-1, 4A-2, and Δ264 tumors. (**G**) Venn diagram showing the relationship of genes significantly (FDR < 0.05) increased in the 4A-1, 4A-2, and Δ264 tumors. (**H**) GO enrichment analysis of genes with significantly (FDR < 0.05) decreased expression in the 4A-1, 4A-2, and Δ264 tumors. (**I**) GO enrichment analysis of genes significantly (FDR < 0.05) increased in the 4A-1, 4A-2, and Δ264 tumors.

## REFERENCES

Adams, C.M., Kim, A.S., Mitra, R., Choi, J.K., Gong, J.Z., and Eischen, C.M. (2017). BCL-W has a fundamental role in B cell survival and lymphomagenesis. J Clin Invest 127, 635–650.

Alimova, I., Pierce, A., Danis, E., Donson, A., Birks, D.K., Griesinger, A., Foreman, N.K., Santi, M., Soucek, L., Venkataraman, S., et al. (2019). Inhibition of MYC attenuates tumor cell self-renewal and promotes senescence in SMARCB1-deficient Group 2 atypical teratoid rhabdoid tumors to suppress tumor growth in vivo. Int J Cancer 144, 1983–1995.

Baluapuri, A., Wolf, E., and Eilers, M. (2020). Target gene-independent functions of MYC oncoproteins. Nat Rev Mol Cell Biol.

Beaulieu, M.E., Jauset, T., Masso-Valles, D., Martinez-Martin, S., Rahl, P., Maltais, L., Zacarias-Fluck, M.F., Casacuberta-Serra, S., Serrano Del Pozo, E., Fiore, C., et al. (2019). Intrinsic cell-penetrating activity propels Omomyc from proof of concept to viable anti-MYC therapy. Sci Transl Med 11.

Bemark, M., and Neuberger, M.S. (2000). The c-MYC allele that is translocated into the IgH locus undergoes constitutive hypermutation in a Burkitt’s lymphoma line. Oncogene 19, 3404–3410.

Blackwell, T.K., Huang, J., Ma, A., Kretzner, L., Alt, F.W., Eisenman, R.N., and Weintraub, H. (1993). Binding of myc proteins to canonical and noncanonical DNA sequences. Mol Cell Biol 13, 5216–5224.

Blackwood, E.M., and Eisenman, R.N. (1991). Max: a helix-loop-helix zipper protein that forms a sequence-specific DNA-binding complex with Myc. Science 251, 1211–1217.

Brockmann, M., Poon, E., Berry, T., Carstensen, A., Deubzer, H.E., Rycak, L., Jamin, Y., Thway, K., Robinson, S.P., Roels, F., et al. (2013). Small molecule inhibitors of aurora-a induce proteasomal degradation of N-myc in childhood neuroblastoma. Cancer Cell 24, 75–89.

Bryan, A.F., Wang, J., Howard, G.C., Guarnaccia, A.D., Woodley, C.M., Aho, E.R., Rellinger, E.J., Matlock, B.K., Flaherty, D.K., Lorey, S.L., et al. (2020). WDR5 is a conserved regulator of protein synthesis gene expression. Nucleic Acids Res 48, 2924–2941.

Cai, Y., Jin, J., Swanson, S.K., Cole, M.D., Choi, S.H., Florens, L., Washburn, M.P., Conaway, J.W., and Conaway, R.C. (2010). Subunit composition and substrate specificity of a MOF-containing histone acetyltransferase distinct from the male-specific lethal (MSL) complex. J Biol Chem 285, 4268–4272.

Capotosti, F., Guernier, S., Lammers, F., Waridel, P., Cai, Y., Jin, J., Conaway, J.W., Conaway, R.C., and Herr, W. (2011). O-GlcNAc transferase catalyzes site-specific proteolysis of HCF-1. Cell 144, 376–388.

Chong, J., Wishart, D.S., and Xia, J. (2019). Using MetaboAnalyst 4.0 for Comprehensive and Integrative Metabolomics Data Analysis. Curr Protoc Bioinformatics 68, e86.

Chou, T.Y., Dang, C.V., and Hart, G.W. (1995). Glycosylation of the c-Myc transactivation domain. Proceedings of the National Academy of Sciences of the United States of America 92, 4417–4421.

Cong, L., Ran, F.A., Cox, D., Lin, S., Barretto, R., Habib, N., Hsu, P.D., Wu, X., Jiang, W., Marraffini, L.A., et al. (2013). Multiplex genome engineering using CRISPR/Cas systems. Science 339, 819–823.

Cowling, V.H., Chandriani, S., Whitfield, M.L., and Cole, M.D. (2006). A Conserved Myc Protein Domain, MBIV, Regulates DNA Binding, Apoptosis, Transformation, and G2 Arrest. Molecular and Cellular Biology 26, 4226–4239.

Dang, C.V., Reddy, E.P., Shokat, K.M., and Soucek, L. (2017). Drugging the ‘undruggable’ cancer targets. Nat Rev Cancer 17, 502–508.

Dingar, D., Kalkat, M., Chan, P.K., Srikumar, T., Bailey, S.D., Tu, W.B., Coyaud, E., Ponzielli, R., Kolyar, M., Jurisica, I., et al. (2014). BioID identifies novel c-MYC interacting partners in cultured cells and xenograft tumors. Journal of proteomics.

Dobin, A., Davis, C.A., Schlesinger, F., Drenkow, J., Zaleski, C., Jha, S., Batut, P., Chaisson, M., and Gingeras, T.R. (2013). STAR: ultrafast universal RNA-seq aligner. Bioinformatics 29, 15–21.

Fahy, E., Subramaniam, S., Murphy, R.C., Nishijima, M., Raetz, C.R., Shimizu, T., Spener, F., van Meer, G., Wakelam, M.J., and Dennis, E.A. (2009). Update of the LIPID MAPS comprehensive classification system for lipids. J Lipid Res 50 Suppl, S9–14.

Farina, A., Faiola, F., and Martinez, E. (2004). Reconstitution of an E box-binding Myc:Max complex with recombinant full-length proteins expressed in Escherichia coli. Protein Expr Purif 34, 215–222.

Feng, J., Liu, T., Qin, B., Zhang, Y., and Liu, X.S. (2012). Identifying ChIP-seq enrichment using MACS. Nat Protoc 7, 1728–1740.

Freiman, R.N., and Herr, W. (1997). Viral mimicry: Common mode of association with HCF by VP16 and the cellular protein LZIP. Genes and Development 11, 3122–3127.

Furrer, M., Balbi, M., Albarca-Aguilera, M., Gallant, M., Herr, W., and Gallant, P. (2010). Drosophila Myc interacts with host cell factor (dHCF) to activate transcription and control growth. Journal of Biological Chemistry 285, 39623–39636.

Giuriato, S., Ryeom, S., Fan, A.C., Bachireddy, P., Lynch, R.C., Rioth, M.J., van Riggelen, J., Kopelman, A.M., Passegue, E., Tang, F., et al. (2006). Sustained regression of tumors upon MYC inactivation requires p53 or thrombospondin-1 to reverse the angiogenic switch. Proc Natl Acad Sci U S A 103, 16266–16271.

Glinsky, G.V., Berezovska, O., and Glinskii, A.B. (2005). Microarray analysis identifies a death-from-cancer signature predicting therapy failure in patients with multiple types of cancer. J Clin Invest 115, 1503–1521.

Goto, H., Motomura, S., Wilson, A.C., Freiman, R.N., Nakabeppu, Y., Fukushima, K., Fujishima, M., Herr, W., and Nishimoto, T. (1997). A single-point mutation in HCF causes temperature-sensitive cell-cycle arrest and disrupts VP16 function. Genes Dev 11, 726–737.

Guan, B.J., Krokowski, D., Majumder, M., Schmotzer, C.L., Kimball, S.R., Merrick, W.C., Koromilas, A.E., and Hatzoglou, M. (2014). Translational control during endoplasmic reticulum stress beyond phosphorylation of the translation initiation factor eIF2alpha. J Biol Chem 289, 12593–12611.

Harding, H.P., Novoa, I., Zhang, Y., Zeng, H., Wek, R., Schapira, M., and Ron, D. (2000). Regulated translation initiation controls stress-induced gene expression in mammalian cells. Mol Cell 6, 1099–1108.

Herbst, A., Hemann, M.T., Tworkowski, K.A., Salghetti, S.E., Lowe, S.W., and Tansey, W.P. (2005). A conserved element in Myc that negatively regulates its proapoptotic activity. EMBO Rep 6, 177–183.

Hershey, J.W. (1991). Translational control in mammalian cells. Annu Rev Biochem 60, 717–755.

Hosios, A.M., Hecht, V.C., Danai, L.V., Johnson, M.O., Rathmell, J.C., Steinhauser, M.L., Manalis, S.R., and Vander Heiden, M.G. (2016). Amino Acids Rather than Glucose Account for the Majority of Cell Mass in Proliferating Mammalian Cells. Dev Cell 36, 540–549.

Iritani, B.M., and Eisenman, R.N. (1999). c-Myc enhances protein synthesis and cell size during B lymphocyte development. Proc Natl Acad Sci U S A 96, 13180–13185.

Itkonen, H.M., Urbanucci, A., Martin, S.E., Khan, A., Mathelier, A., Thiede, B., Walker, S., and Mills, I.G. (2019). High OGT activity is essential for MYC-driven proliferation of prostate cancer cells. Theranostics 9, 2183–2197.

Iwata, T.N., Cowley, T.J., Sloma, M., Ji, Y., Kim, H., Qi, L., and Lee, S.S. (2013). The transcriptional co-regulator HCF-1 is required for INS-1 β-cell glucose-stimulated insulin secretion. PLoS One 8, e78841.

Jain, M., Arvanitis, C., Chu, K., Dewey, W., Leonhardt, E., Trinh, M., Sundberg, C.D., Bishop, J.M., and Felsher, D.W. (2002). Sustained loss of a neoplastic phenotype by brief inactivation of MYC. Science 297, 102–104.

Julien, E., and Herr, W. (2003). Proteolytic processing is necessary to separate and ensure proper cell growth and cytokinesis functions of HCF-1. EMBO Journal 22, 2360–2369.

Kalkat, M., Resetca, D., Lourenco, C., Chan, P.K., Wei, Y., Shiah, Y.J., Vitkin, N., Tong, Y., Sunnerhagen, M., Done, S.J., et al. (2018). MYC Protein Interactome Profiling Reveals Functionally Distinct Regions that Cooperate to Drive Tumorigenesis. Mol Cell 72, 836–848 e837.

Kalkat, M., Wasylishen, A.R., Kim, S.S., and Penn, L. (2011). More than MAX: Discovering the Myc interactome. Cell Cycle 10, 374–375.

Kent, L.N., and Leone, G. (2019). The broken cycle: E2F dysfunction in cancer. Nat Rev Cancer 19, 326–338.

Klein, G., Giovanella, B., Westman, A., Stehlin, J.S., and Mumford, D. (1975). An EBV-genome-negative cell line established from an American Burkitt lymphoma; receptor characteristics. EBV infectibility and permanent conversion into EBV-positive sublines by in vitro infection. Intervirology 5, 319–334.

Krzywinski, M., Schein, J., Birol, I., Connors, J., Gascoyne, R., Horsman, D., Jones, S.J., and Marra, M.A. (2009). Circos: an information aesthetic for comparative genomics. Genome Res 19, 1639–1645.

Kurland, J.F., and Tansey, W.P. (2008). Myc-mediated transcriptional repression by recruitment of histone deacetylase. Cancer Res 68, 3624–3629.

Laferte, A., Favry, E., Sentenac, A., Riva, M., Carles, C., and Chedin, S. (2006). The transcriptional activity of RNA polymerase I is a key determinant for the level of all ribosome components. Genes Dev 20, 2030–2040.

Langmead, B., Trapnell, C., Pop, M., and Salzberg, S.L. (2009). Ultrafast and memory-efficient alignment of short DNA sequences to the human genome. Genome Biol 10, R25.

Levens, D. (2002). Disentangling the MYC web. Proc Natl Acad Sci U S A 99, 5757–5759.

Liao, Y., Smyth, G.K., and Shi, W. (2014). featureCounts: an efficient general purpose program for assigning sequence reads to genomic features. Bioinformatics 30, 923–930.

Love, M.I., Huber, W., and Anders, S. (2014). Moderated estimation of fold change and dispersion for RNA-seq data with DESeq2. Genome Biol 15, 550.

Machida, Y.J., Machida, Y., Vashisht, A.A., Wohlschlegel, J.A., and Dutta, A. (2009). The deubiquitinating enzyme BAP1 regulates cell growth via interaction with HCF-1. J Biol Chem 284, 34179–34188.

Martin, M. (2011). Cutadapt removes adapter sequences from high-throughput sequencing reads. EMBnet journal 17, 10–12.

Mazars, R., Gonzalez-de-Peredo, A., Cayrol, C., Lavigne, A.C., Vogel, J.L., Ortega, N., Lacroix, C., Gautier, V., Huet, G., Ray, A., et al. (2010). The THAP-zinc finger protein THAP1 associates with coactivator HCF-1 and O-GlcNAc transferase: a link between DYT6 and DYT3 dystonias. J Biol Chem 285, 13364–13371.

McMahon, S.B., Van Buskirk, H.A., Dugan, K.A., Copeland, T.D., and Cole, M.D. (1998). The novel ATM-related protein TRRAP is an essential cofactor for the c-Myc and E2F oncoproteins. Cell 94, 363–374.

Meyer, N., and Penn, L.Z. (2008). Reflecting on 25 years with MYC. Nature reviews Cancer 8, 976–990.

Miller, S.A., Dykes, D.D., and Polesky, H.F. (1988). A simple salting out procedure for extracting DNA from human nucleated cells. Nucleic Acids Res 16, 1215.

Minocha, S., Villeneuve, D., Praz, V., Moret, C., Lopes, M., Pinatel, D., Rib, L., Guex, N., and Herr, W. (2019). Rapid Recapitulation of Nonalcoholic Steatohepatitis upon Loss of Host Cell Factor 1 Function in Mouse Hepatocytes. Mol Cell Biol 39.

Morrish, F., Giedt, C., and Hockenbery, D. (2003). c-MYC apoptotic function is mediated by NRF-1 target genes. Genes Dev 17, 240–255.

Morrish, F., and Hockenbery, D. (2014). MYC and mitochondrial biogenesis. Cold Spring Harb Perspect Med 4.

Nabet, B., Roberts, J.M., Buckley, D.L., Paulk, J., Dastjerdi, S., Yang, A., Leggett, A.L., Erb, M.A., Lawlor, M.A., Souza, A., et al. (2018). The dTAG system for immediate and target-specific protein degradation. Nat Chem Biol 14, 431–441.

Nie, Z., Hu, G., Wei, G., Cui, K., Yamane, A., Resch, W., Wang, R., Green, D.R., Tessarollo, L., Casellas, R., et al. (2012). c-Myc is a universal amplifier of expressed genes in lymphocytes and embryonic stem cells. Cell 151, 68–79.

Parker, J.B., Yin, H., Vinckevicius, A., and Chakravarti, D. (2014). Host cell factor-1 recruitment to E2F-bound and cell-cycle-control genes is mediated by THAP11 and ZNF143. Cell Rep 9, 967–982.

Salvat, F., Jablonski, A., Powell, C.J., and Lee, A.Y. (2016). NIST Electron Elastic-Scattering Cross-Section Database, Version 4.0 (Gaithersburg, MD: National Institute of Standards and Technology).

Schrimpe-Rutledge, A.C., Codreanu, S.G., Sherrod, S.D., and McLean, J.A. (2016). Untargeted Metabolomics Strategies-Challenges and Emerging Directions. J Am Soc Mass Spectrom 27, 1897–1905.

Scott, M., Klumpp, S., Mateescu, E.M., and Hwa, T. (2014). Emergence of robust growth laws from optimal regulation of ribosome synthesis. Mol Syst Biol 10, 747.

Shannon, P., Markiel, A., Ozier, O., Baliga, N.S., Wang, J.T., Ramage, D., Amin, N., Schwikowski, B., and Ideker, T. (2003). Cytoscape: a software environment for integrated models of biomolecular interaction networks. Genome Res 13, 2498–2504.

Smith, C.A., O’Maille, G., Want, E.J., Qin, C., Trauger, S.A., Brandon, T.R., Custodio, D.E., Abagyan, R., and Siuzdak, G. (2005). METLIN: a metabolite mass spectral database. Ther Drug Monit 27, 747–751.

Soucek, L., Whitfield, J.R., Sodir, N.M., Masso-Valles, D., Serrano, E., Karnezis, A.N., Swigart, L.B., and Evan, G.I. (2013). Inhibition of Myc family proteins eradicates KRas-driven lung cancer in mice. Genes Dev 27, 504–513.

Southern, E.M. (1975). Detection of specific sequences among DNA fragments separated by gel electrophoresis. J Mol Biol 98, 503–517.

Stark, R., and Brown, G. (2011). DiffBind: differential binding analysis of ChIP-Seq peak data. Bioconductor.

Swamy, M., Pathak, S., Grzes, K.M., Damerow, S., Sinclair, L.V., van Aalten, D.M., and Cantrell, D.A. (2016). Glucose and glutamine fuel protein O-GlcNAcylation to control T cell self-renewal and malignancy. Nat Immunol 17, 712–720.

Tansey, W.P. (2014). Mammalian MYC Proteins and Cancer. New Journal of Science 2014, 1–27.

Thomas, L.R., Adams, C.M., Wang, J., Weissmiller, A.M., Creighton, J., Lorey, S.L., Liu, Q., Fesik, S.W., Eischen, C.M., and Tansey, W.P. (2019). Interaction of the oncoprotein transcription factor MYC with its chromatin cofactor WDR5 is essential for tumor maintenance. Proc Natl Acad Sci U S A 116, 25260–25268.

Thomas, L.R., Foshage, A.M., Weissmiller, A.M., Popay, T.M., Grieb, B.C., Qualls, S.J., Ng, V., Carboneau, B., Lorey, S., Eischen, C.M., et al. (2016). Interaction of MYC with host cell factor-1 is mediated by the evolutionarily conserved Myc box IV motif. Oncogene 35, 3613–3618.

Thomas, L.R., Wang, Q., Grieb, B.C., Phan, J., Foshage, A.M., Sun, Q., Olejniczak, E.T., Clark, T., Dey, S., Lorey, S., et al. (2015). Interaction with WDR5 promotes target gene recognition and tumorigenesis by MYC. Mol Cell 58, 440–452.

Tyagi, S., Chabes, A.L., Wysocka, J., and Herr, W. (2007). E2F activation of S phase promoters via association with HCF-1 and the MLL family of histone H3K4 methyltransferases. Mol Cell 27, 107–119.

van Riggelen, J., Yetil, A., and Felsher, D.W. (2010). MYC as a regulator of ribosome biogenesis and protein synthesis. Nat Rev Cancer 10, 301–309.

Venkateswaran, N., Lafita-Navarro, M.C., Hao, Y.H., Kilgore, J.A., Perez-Castro, L., Braverman, J., Borenstein-Auerbach, N., Kim, M., Lesner, N.P., Mishra, P., et al. (2019). MYC promotes tryptophan uptake and metabolism by the kynurenine pathway in colon cancer. Genes Dev 33, 1236–1251.

Welcker, M., Orian, A., Jin, J., Grim, J.A., Harper, J.W., Eisenman, R.N., and Clurman, B.E. (2004). The Fbw7 tumor suppressor regulates glycogen synthase kinase 3 phosphorylation-dependent c-Myc protein degradation. Proc Natl Acad Sci U S A 101, 9085–9090.

Wickham, H. (2016). ggplot2: Elegant Graphics for Data Analysis (Springer International Publishing).

Wiman, K.G., Clarkson, B., Hayday, A.C., Saito, H., Tonegawa, S., and Hayward, W.S. (1984). Activation of a translocated c-myc gene: role of structural alterations in the upstream region. Proc Natl Acad Sci U S A 81, 6798–6802.

Wise, D.R., DeBerardinis, R.J., Mancuso, A., Sayed, N., Zhang, X.Y., Pfeiffer, H.K., Nissim, I., Daikhin, E., Yudkoff, M., McMahon, S.B., et al. (2008). Myc regulates a transcriptional program that stimulates mitochondrial glutaminolysis and leads to glutamine addiction. Proc Natl Acad Sci U S A 105, 18782–18787.

Wishart, D.S., Feunang, Y.D., Marcu, A., Guo, A.C., Liang, K., Vazquez-Fresno, R., Sajed, T., Johnson, D., Li, C., Karu, N., et al. (2018). HMDB 4.0: the human metabolome database for 2018. Nucleic Acids Res 46, D608–D617.

Wishart, D.S., Jewison, T., Guo, A.C., Wilson, M., Knox, C., Liu, Y., Djoumbou, Y., Mandal, R., Aziat, F., Dong, E., et al. (2013). HMDB 3.0--The Human Metabolome Database in 2013. Nucleic Acids Res 41, D801–807.

Wysocka, J., and Herr, W. (2003). The herpes simplex virus VP16-induced complex: the makings of a regulatory switch. Trends Biochem Sci 28, 294–304.

Wysocka, J., Myers, M.P., Laherty, C.D., Eisenman, R.N., and Herr, W. (2003). Human Sin3 deacetylase and trithorax-related Set1/Ash2 histone H3-K4 methyltransferase are tethered together selectively by the cell-proliferation factor HCF-1. Genes Dev 17, 896–911.

Zhou, Y., Zhou, B., Pache, L., Chang, M., Khodabakhshi, A.H., Tanaseichuk, O., Benner, C., and Chanda, S.K. (2019). Metascape provides a biologist-oriented resource for the analysis of systems-level datasets. Nat Commun 10, 1523.

